# Systematic identification of a nuclear receptor-enriched predictive signature for erastin-induced ferroptosis

**DOI:** 10.1101/2020.04.13.038430

**Authors:** Ok-Seon Kwon, Eun-Ji Kwon, Hyeon-Joon Kong, Jeong-Yoon Choi, Yun-Jeong Kim, Eun-Woo Lee, Wankyu Kim, Haeseung Lee, Hyuk-Jin Cha

**Affiliations:** Stem Cell Convergence Research Center, Korea Research Institute of Bioscience and Biotechnology (KRIBB), Daejeon 34141, Korea; College of Pharmacy, Seoul National University, Seoul 08826, Republic of Korea; Metabolic Regulation Research Center, Korea Research Institute of Bioscience and Biotechnology (KRIBB), Daejeon 34141, Korea; Department of Life Sciences, Ewha Womans University, Seoul 03760, Republic of Korea; Research Institute of Pharmaceutical Sciences, Seoul National University, Seoul 08826, Republic of Korea

**Author notes:** Corresponding Authors and, Hyuk-Jin Cha, Ph.D., Research institute of pharmaceutical sciences, Seoul National University, 1 Gwanak-ro, Gwanak-gu, Seoul 08826, Republic of Korea, Tel.: ^+^82-2-880-7825; Fax: ^+^82-2-880-9122;, Haeseung Lee, Ph.D. Intellectual Information Team, Future Medicine Division, Korea Institute of Oriental Medicine, Daejeon 34054, Korea, Tel.: ^+^82-42-868-9268.

**Keywords:** Erastin, Ferroptosis, Redox imbalance, NRF2, Aryl hydrocarbon receptor, Elastic net, Drug response biomarker

## Abstract

Erastin, which has been initially identified as a synthetic lethal compound against cancer expressing an RAS oncogene, inhibits cystine/glutamate antiporters and causes ferroptic cell death in various cell types, including therapy-resistant mesenchymal cancer cells. However, despite recent emerging evidence for the mechanisms underlying ferroptosis, molecular biomarkers associated with erastin-dependent ferroptosis have not yet been identified. In the present study, we employed isogenic lung cancer cell models with therapy-resistant mesenchymal properties to show that a redox imbalance leads to glutathione depletion and ferroptotic cell death. Subsequent gene expression analysis of pan-cancer cell lines revealed that the activity of transcription factors, including nuclear factor erythroid 2-related factor 2 (NRF2) and aryl hydrocarbon receptor (AhR), serve as important markers of erastin resistance. Based on the integrated expression of genes in the nuclear receptor meta-pathway (NRM), we constructed an NRM model and validated its robustness using an independent pharmacogenomics dataset. The NRM model was further evaluated by employing it in the sensitivity testing of nine cancer cell lines for which erastin sensitivities had not yet been undetermined. Our pharmacogenomics approach has the potential to pave the way for the efficient classification of patients for therapeutic intervention using erastin or erastin analogs.

## Introduction

Erastin (which derives its name from being an eradicator of RAS and ST-expressing cells) is a small molecule that was first reported as inducing synthetic lethality in cancer cells expressing an RAS oncogene [1] through an oxidative stress mechanism under strong RAS-RAF-MEK signaling [2]. The mode of action (MoA) for cell death was subsequently identified as ferroptosis, a unique iron-dependent form of nonapoptotic cell death [3]. Erastin inhibits system X_c_^-^ (XCT), thus impairing the cystine/glutamate antiporter (encoded by *SLC7A11*) that is involved in the synthesis of glutathione (GSH) from imported cystine and creating a void in antioxidant defense that leads to ferroptosis [3]. Small molecule inhibitors of ferroptosis have since been developed for a variety of therapeutic applications to inhibit pathological cell death, including the treatment of neurodegenerative diseases, stroke, and ischemic injuries [4]. In particular, ferroptosis inducers (FINs) such as erastin have been extensively examined as novel anti-cancer therapeutics [5, 6]. However, to date, clinical trials of FINs such as sulfasalazine, which is an inhibitor of XCT [7] in glioma patients, have been unsatisfactory due to the lack of clinical response [8].

The high dependency of therapy-resistant mesenchymal cancer cells (with high *ZEB1* expression) on the lipid peroxidase pathway governed by phospholipid glutathione peroxidase 4 (*GPX4*) increases their vulnerability to ferroptosis via *GPX4* inhibition or GSH depletion with erastin treatment [9]. *GPX4* dependency on erastin induced ferroptic cell death occurs in cell-type-specific manner [10], though the strength of this vulnerability varies in a cell-specific manner. As such, a variety of cellular and molecular components and processes, such as metabolic heterogeneity[11], mesenchymal properties [9], differentiation status [12], p53 status [13], transcription factors [14, 15], signaling pathways (e.g. MAPK[16], ATM [17] or YAP [18]), integrins [19], GSH regulators [20], and levels of monounsaturated fatty acid [21], have been examined as determinants of ferroptosis vulnerability in a diverse range of cell model systems. Despite this, the variation in the susceptibility of cancer cells to ferroptosis, via either XCT or GPX4 inhibition, depending on cellular and molecular characteristics has not yet been fully understood. In this regard, the establishment of a unique signature that enabled the prediction of erastin vulnerability would be useful for patient stratification, which would maximize the efficacy and minimize the toxicity of anti-cancer therapy using erastin analogs that are currently being tested in clinical trials [22].

The pharmacogenomics approach has advanced the understanding of the MoA of various drugs by systematically identifying molecular biomarkers that contribute to drug responses [23, 24]. In this respect, gene expression data have been found to be the most informative of available omics datasets (e.g., genomic, proteomic, and epigenomic profiling data) in predicting the drug response of human cancer cells [25, 26]. In precision oncology, transcriptomic profiling has been widely employed to screen for predictive gene signatures that effectively guide treatment decisions using a few to a thousand cultured cell lines as surrogates [9, 12, 27, 28]. The key resources behind these efforts are the Cancer Cell Line Encyclopedia (CCLE) and the Cancer Therapeutics Response Portal (CTRP); these databases provide both transcriptomic data and data from the sensitivity screening of 860 cancer cell lines against 487 compounds [29, 30]. These datasets make it possible to revisit the MoA of particular drugs by offering robust molecular signatures from distinct features of cell lines that exhibit differences in their drug sensitivity.

In the present study, we constructed an effective model for the prediction of erastin sensitivity based on the basal gene expression and drug-response profiles of pan-cancer cell lines obtained from the CCLE and CTRP datasets. This model revealed that nuclear receptor-enriched gene signatures are important determinants of erastin-induced ferroptotic cell death. Our approach accurately predicts the erastin sensitivity of cancer cell lines based on their basal gene expression, indicating that it would be useful for identifying patients who could potentially respond to erastin treatment.

## Materials and Methods

### RNA sequencing (RNA-seq) and data processing

Total RNA was isolated using Trizol according to the manufacturer’s instructions. For library construction, we used the TruSeq Stranded mRNA Library Prep Kit (Illumina, San Diego, CA). Briefly, the strand-specific protocol included the following steps: (1) strand cDNA synthesis, (2) strand synthesis using dUTPs instead of dTTPs, (3) end repair, A-tailing, and adaptor ligation, and (4) PCR amplification. Each library was then diluted to 8 pM for 76 cycles of paired-read sequencing (2 × 75 bp) on an Illumina NextSeq 500 following the manufacturer’s recommended protocol.

The sequencing quality of the raw FASTQ files was assessed using FastQC (https://www.bioinformatics.babraham.ac.uk/projects/fastqc/). Low-quality reads and the adapter sequences within these reads were eliminated using BBDuk (http://jgi.doe.gov/data-and-tools/bb-tools/). Trimmed reads were aligned to the GRCh37 reference genome (build 38) using the STAR aligner (v2.6.0a). Gene-level transcripts per million (TPM) and read counts were calculated using RSEM v.1.3.1. with Gencode v19 annotation. The FASTQ files and processed data are available in the Gene Expression Omnibus (GEO: GSE135402). Genes differentially expressed between A549 and TD cells were obtained using the DESeq2 package in R.

### Cancer cell line RNA-seq and erastin sensitivity data

Baseline gene-expression profiles of 932 cancer cell lines were downloaded from the NCI’s Genomic Data Commons (GDC, https://gdc.cancer.gov/) as BAM files. Gene-level TPM and expected counts were quantified using RSEM v.1.3.1. with Gencode v19 annotation. A total of 18,965 protein-coding genes were retained for model training and subsequent analysis. The erastin drug-response profiles of 804 cancer cell lines were obtained from CTD^2^ Data Portal (https://ocg.cancer.gov/programs/ctd2/data-portal). Cell viability data were converted into growth inhibition data and adjusted to fall within a range of 0–100 %. The adjusted growth inhibition data were subjected to four-parameter logistic regression and low-quality profiles (goodness of fit < 0.7) removed. The dose-response area under the curve (AUC) for sensitivity was normalized to a range of 0–1 using the maximum AUC, which was assumed to represent 0 % growth inhibition, for a given concentration range.

### Predictive modeling of erastin sensitivity

To identify the genes that were most predictive of erastin sensitivity, we adapted an elastic net regression approach, which is a penalized model widely used for feature selection, particularly with genome-scale data [31]. A total of 598 non-hematologic cancer cell lines with available RNA-seq (TPM) and erastin sensitivity (AUC) data were used to build multiple models, each of which considered the expression of genes within the top 16 individually enriched pathways as a feature set (i.e., one model per pathway). Each model was assessed using nested leave-one-out cross-validation (LOOCV) in which a single sample (i.e., a cell line) was used to test a model trained by the remaining samples (i.e., the other 597 cell lines). This process was repeated until all cell lines had been used as the test dataset. For each training run, the optimal parameters (α and λ) were taken to be those that minimized the mean square error for five-fold cross-validation with 10 iterations of the training data. Predicted AUC values from each test set were concatenated and then compared to the actual AUC data using Spearman correlation to evaluate prediction performance. The same procedure was also applied to assess the generalized linear regression models in Figure 4D. For interpretation and visualization purposes, the predicted AUCs were scaled to the distribution of the actual AUC data. The overall process was conducted using the glmnet and caret packages in R.

**Figure 1.**
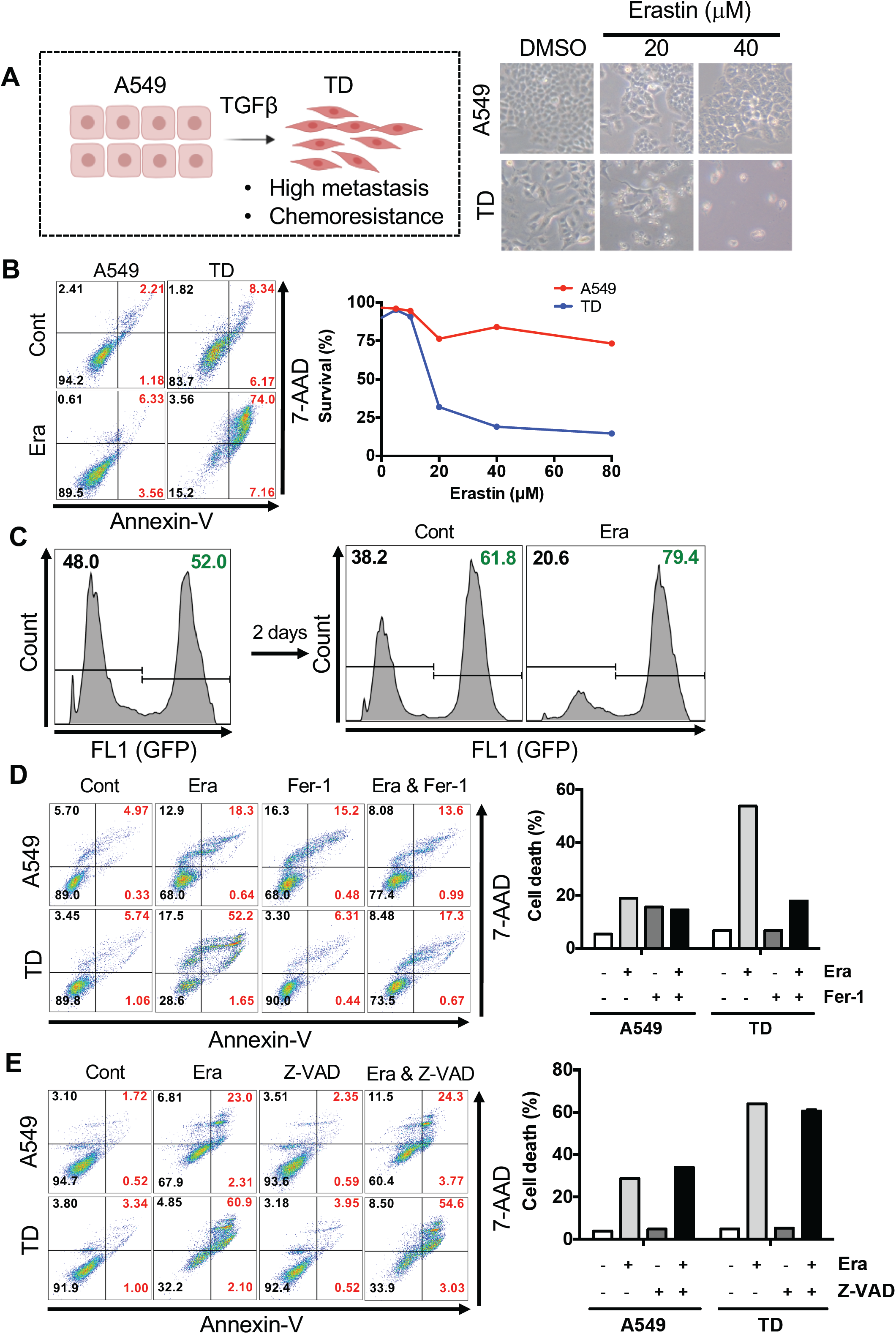
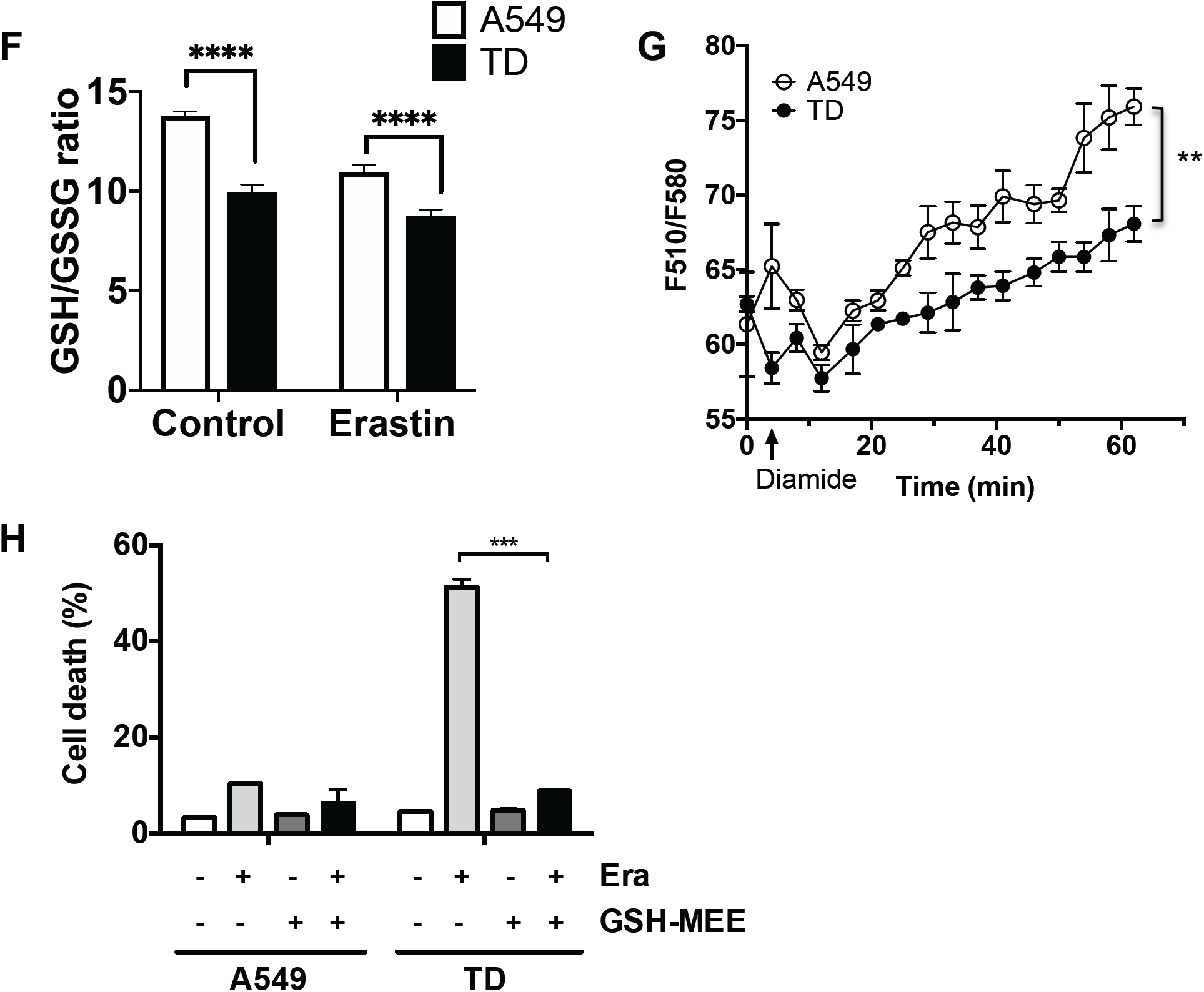
Higher sensitivity to ferroptosis in mesenchymal lung cancer cells. **(A)** Schematic diagram of model for A549 and TD (left). Microscopic images of A549 and TD cells 24 hours after an indicative dose of erastin. **(B)** Flow cytometry for Annexin V and 7-AAD 24 hours after 40 μM erastin treatment. **(C)** Flow cytometry for GFP-positive cells (A549-GFP) compared to GFP-negative cells (TD) two days after 40 μM erastin treatment. **(D, E)** Flow cytometry for Annexin V and 7-AAD 24 hours after 40 μM erastin treatment with or without ferrostatin (Fer-1) pretreatment (D) or Z-VAD (E). **(F)** Fold ratio of GSH/GSSG 24 hours after treatment with 40 μM erastin in A549 or TD cells. **(G)** Relative ratio of the fluorescent intensity of FreSHtracer after treatment with diamide in A549 (open circle) or TD (closed circle) cells at the indicated times. **(H)** Levels of cell death population of A549/TD cells after erastin treatment with or without GSH-MEE.

**Figure 2.**
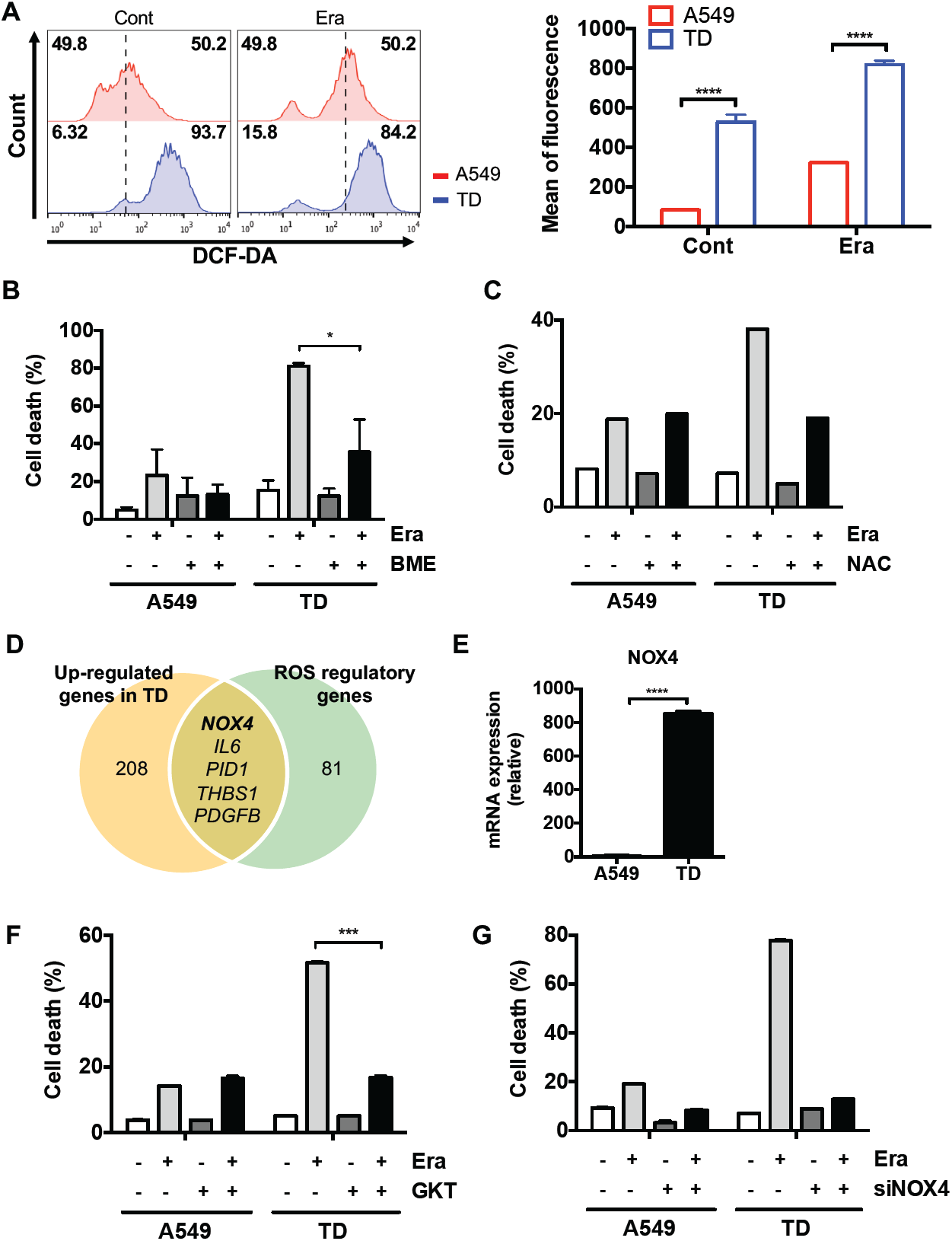
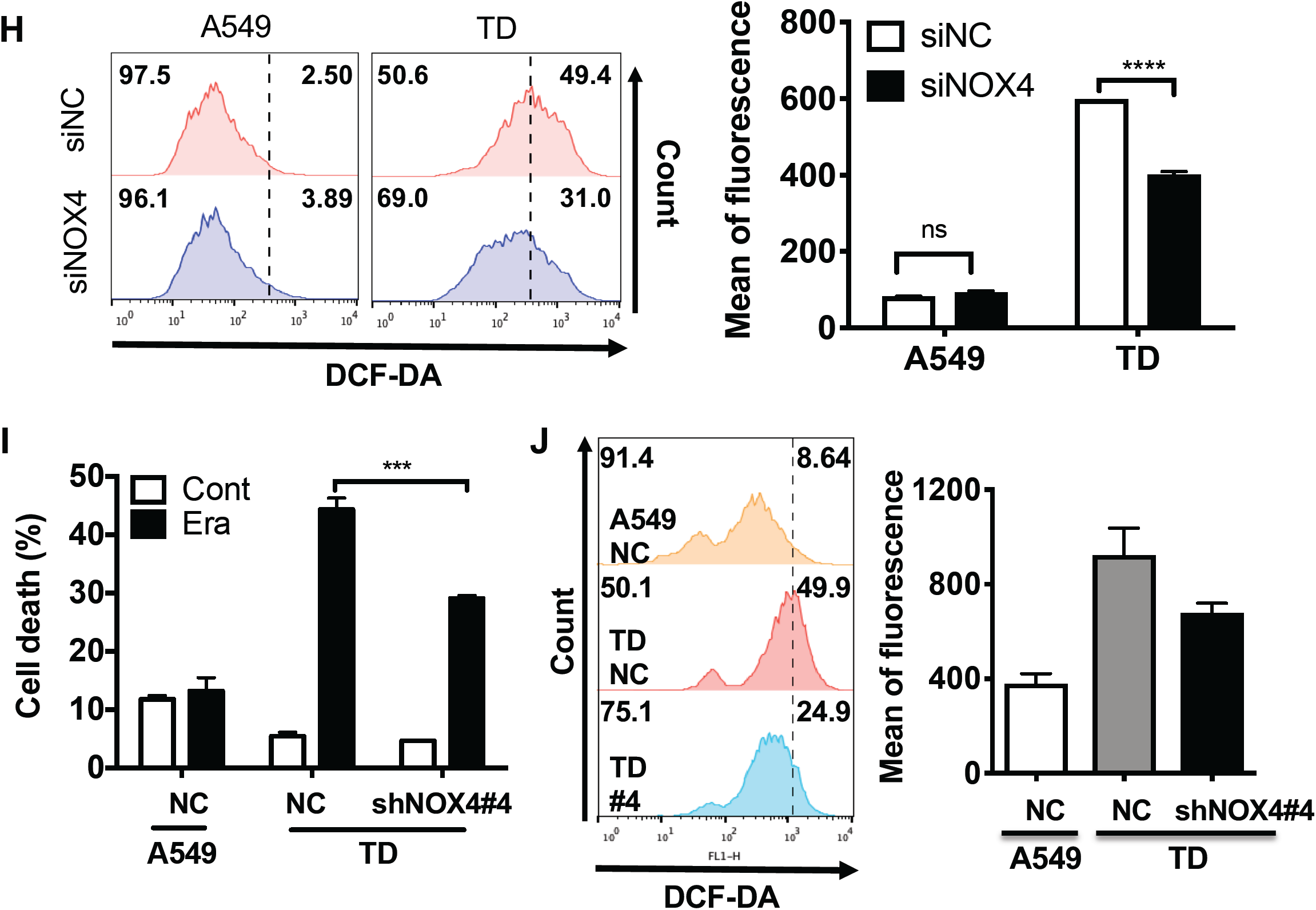
Expenditure of GSH due to the redox imbalance caused by NOX4 induction. **(A)** Flow cytometry for DCF-DA 24 hours after treatment with the vehicle (Cont) or erastin (Era) in A549 or TD cells (left). Graphical representation of the mean fluorescence intensity of DCF-DA (right). **(B, C)** Cell death population size of A549 or TD cells 24 hours after 40 μM erastin treatment with BME (B) or NAC (C) pretreatment. **(D)** Venn diagram showing the number of shared genes involved in ROS regulation (GO: 0000302) and upregulated in TD compared to A549 cells. Upregulated genes that met the criteria of log_2_FC ≥ 4 and FDR ≤ 10^−3^ were selected using DESeq2. **(E)** mRNA expression of NOX4 in A549 and TD cells. **(F, G)** Cell death population size in A549 or TD cells 24 hours after 40 μM erastin treatment following pretreatment with GKT (F) or NOX4 siRNA transfection (G). **(H)** Flow cytometry for DCF-DA 24 hours after the introduction of siRNA for the control (siNC) or NOX4 (siNOX4) in A549 or TD cells (left). Graphical representation of the mean fluorescence intensity of DCF-DA (right). **(I)** Graphical representation of the cell death population 24 hours after 40 μM erastin treatment in the control (NC) or NOX4 knockdown cells (clone #4). **(J)** Flow cytometry for DCF-DA in the A549, TD control (NC), and NOX4-knockdown TD cells (TD#4) (left) and graphical representation of the mean fluorescence intensity of DCF-DA (right).

**Figure 3.**
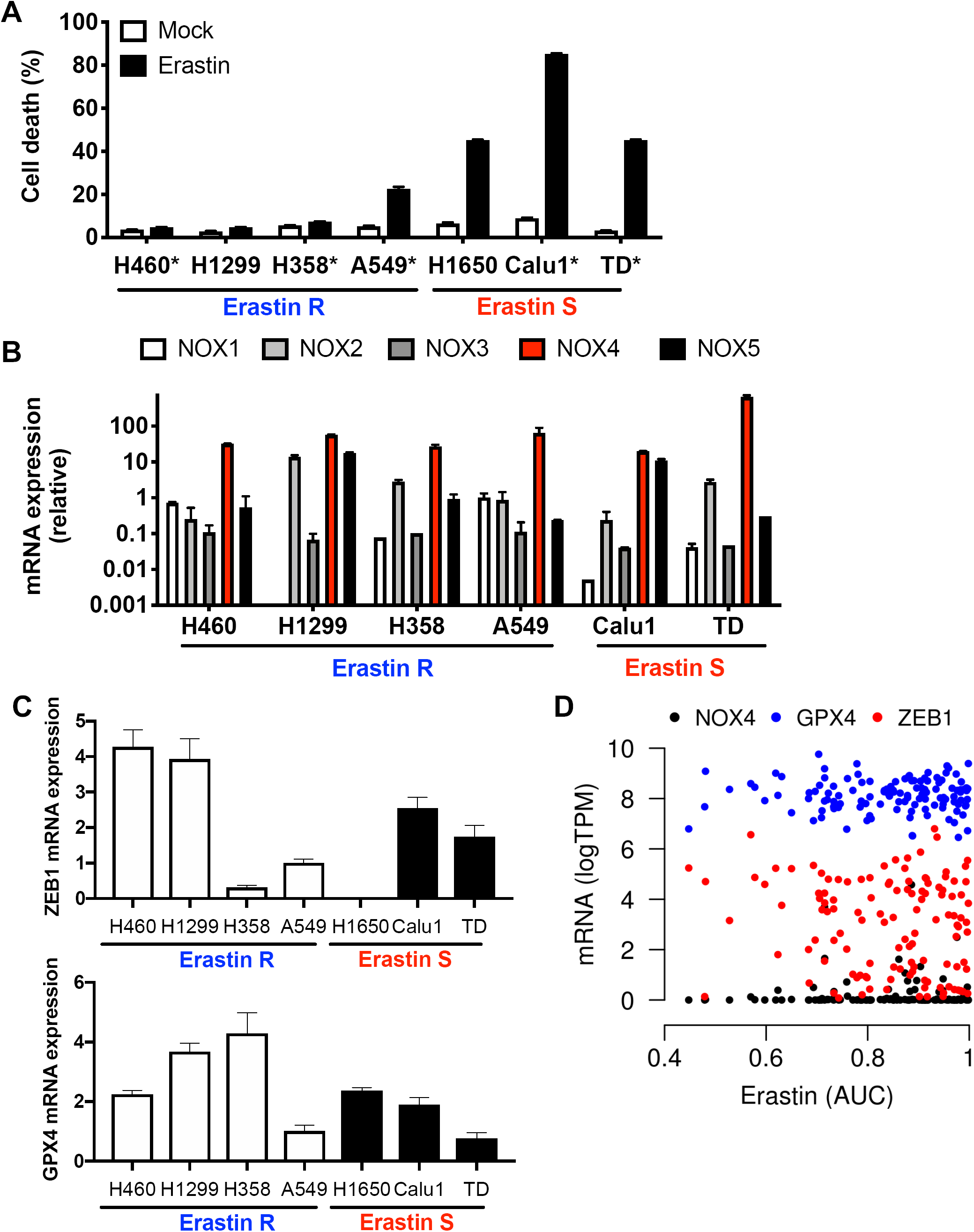
NOX4 as an insufficient marker for erastin sensitivity. **(A)** Percentage of cell death population in the indicated lung cancer cell lines 24 hours after 40 μM erastin treatment (erastin R: resistant, erastin S: sensitive, * *KRAS* mutation). **(B)** mRNA expression of NOXs in the indicated lung cancer cell lines. **(C)** mRNA expression of *ZEB1* (top) and *GPX4* (bottom) in the indicated lung cancer cell lines. **(D)** Relationship between erastin sensitivity and the gene expression of *NOX4, GPX4*, and *ZEB1* in 123 lung cancer cell lines. Erastin sensitivity (AUC) data from the CTRP and gene expression levels from the CCLE RNA-seq (log_2_TPM) are shown.

**Figure 4.**
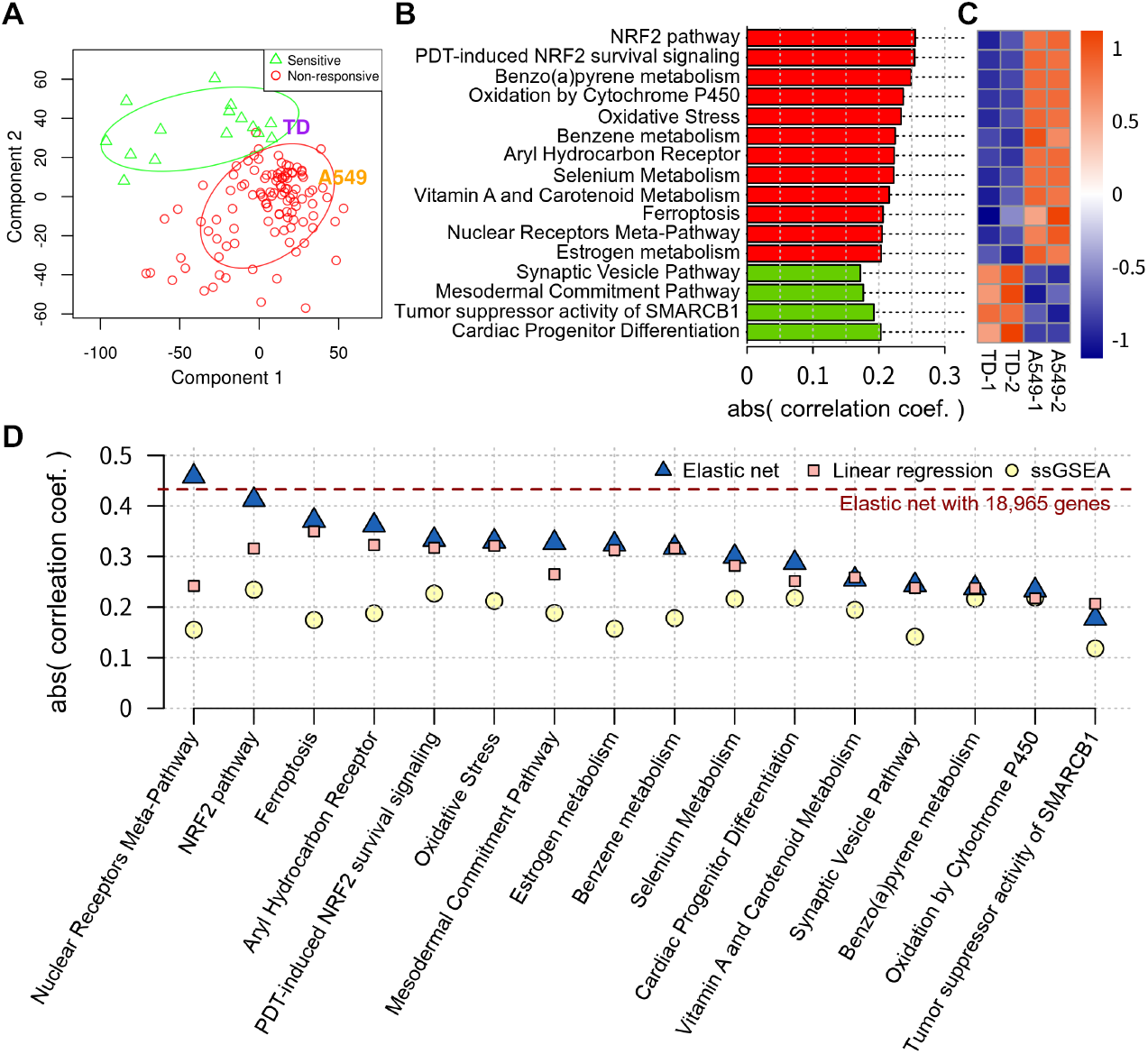
Prediction of the molecular mechanisms that contribute to erastin sensitivity. **(A)** Partial least square discriminant analysis (PLS-DA) of lung cancer cell lines (123 CCLE cell lines, A549, and TD) based on global gene expression profiles. To divide the CCLE cell lines into erastin sensitive (S) and resistant (R) groups, we roughly defined the cut-off (AUC = 0.7) with reference to the AUC values of two sensitive (Calu1, NCI-H1650) and four resistant (NCI-H1299, A549, NCI-H359, NCI-H460) cell lines tested beforehand (Fig. S4A). **(B)** The pathways most closely associated with erastin sensitivity. The association between the pathways and erastin sensitivity was measured using Pearson’s correlation between the cell-line pathway-enrichment scores (PESs) and erastin sensitivity (AUC). Pathways with an absolute *z*-normalized correlation coefficient greater than 2 were selected. Positive and negative correlations are shown in red and green, respectively. Gene annotations for the 445 pathways were obtained from Wikipathways. **(C)** Scaled PESs of the top 16 pathways in A549 and TD cells. **(D)** Performance of individual predictions of erastin sensitivity. The predictions were assessed using leave-one-out cross-validation (LOOCV) with the absolute Spearman’s correlation coefficient for the actual and predicted AUC. Predictions using an elastic net, a general linear model (GLM), and ssGSEA based on each of the 16 feature gene sets are represented. Elastic net predictions employing the expression of all genes (n=18,965) are indicated by the red line.

### Statistical analysis

The statistical significance of any differences among three groups and between two groups was determined using one-way analysis of variance (ANOVA) with multiple comparisons and Student’s *t*-tests (two-tailed), respectively. Significance was set at *P* < 0.05 (*), *P* < 0.01 (**), and *P* < 0.001 (***). The error bars represent the mean ± s.d.

## Results

### Higher sensitivity to ferroptosis in mesenchymal lung cancer cells

In order to examine selective ferroptosis in therapy-resistant mesenchymal cancer cells [9], we used erastin to trigger ferroptosis because a potent analog of erastin, belonging to the group of class I inhibitors targeting XCT is currently undergoing clinical trials for the treatment of cancer [32] due to its *in vivo* suitable pharmacokinetics, a characteristic not shared by other FINs [33]. For the isogenic pairing of epithelial and mesenchymal lung cancer cells, the mesenchymal lung cancer cell lines (transdifferentiated lung cancer cells, hereafter referred to as TD cells) derived from A549 cancer cells following chronic TGFβ exposure [34] were used in this study (Fig. 1A). Consistent with previous reports [27, 35–37], RNA-seq data analysis of the A549 and TD cells (GSE135402) revealed that mesenchymal and therapy-resistant gene signatures were upregulated in TD cells (Fig. S1A). Of particular interest was the fact that TD cells exhibiting chemoresistance [27, 37] were highly sensitive to erastin-induced cell death (Figs. 1A and B). Selective cell death of TD cells after erastin treatment was highlighted when A549 and TD cells were co-cultured (Figs. 1C and S1B and Movie S1). As with the selective death of TD cells, erastin-induced cell death was significantly blocked by ferrostatin-1, a ferroptosis inhibitor (Fig. 1D), but not by a pan-caspase inhibitor (Fig. 1E). We thus concluded that the TD cell death induced by erastin was the result of ferroptosis.

Given the important role of XCT in glutathione (GSH) synthesis, the higher sensitivity of TD cells to erastin (Figs. 1A and B) could be the result of the lower basal levels of reduced GSH compared to oxidized GSH due to the inhibition of XCT by erastin, as previously described [10]. As predicted, the ratio of reduced GSH to oxidized GSH (GSSG) was significantly lower in the TD cells, independent of erastin treatment (Fig. 1F). Thus, we also monitored the levels of reduced GSH in real-time using FreSHtracer, a recently validated real-time fluorescent thiol tracer [38]. Consistent with the results shown in Figure 1F, the recovery of GSH after thiol-specific oxidant diamide treatment was significantly retarded in TD cells when compared to A549 cells (Fig. 1G). Importantly, supplementation with GSH monoethyl ester (GSH-MEE), a cell-permeable derivative of GSH, markedly rescued TD cell death following erastin treatment (Fig. 1H). As a result, it is clear that the lower basal levels of the reduced form of GSH in TD cells cause high sensitivity to erastin-induced ferroptosis, as previously reported [12].

### Expenditure of GSH due to the redox imbalance caused by NOX4 induction

To explain the consistently lower levels of the reduced form of GSH in TD cells, we first examined the reactive oxygen species (ROS) levels in A549 and TD cells. Interestingly, basal ROS levels and ROS levels induced by erastin treatment were much higher in TD cells (Fig. 2A). It is generally accepted that the redox imbalance in cancer cells leads to recurrence, drug resistance, and metastasis [39], which are typical characteristics of the epithelial– mesenchymal transition (EMT). Accordingly, it is assumed that the lower levels of the reduced form of GSH observed in TD cells result from the consistently high levels of ROS. Along the same lines, the anti-oxidants β-mercaptoethanol (BME) and N-acetyl-cystine (NAC) significantly attenuated erastin-induced ferroptosis (Figs. 2B and C).

The redox imbalance that leads to GSH depletion and higher ferroptosis sensitivity in TD cells could be caused by the induction of genes that may affect the mechanisms underlying the regulation of ROS. To identify these genes, we investigated the changes in global gene expression in A549 and TD cells. Of the genes upregulated in TD cells, we focused on ROS regulatory genes (Fig. S2A, red dots) and identified NADPH oxidase 4 (*NOX4*) as a candidate because of the obvious role of NOX4 in both ROS production [40] and ferroptosis [16] (Fig. 2D). High-level *NOX4* expression in TD cells (Fig. 2E) was found to be responsible for erastin-induced ferroptosis because chemical inhibition using GKT-137831 (a NOX1/4 inhibitor) or the knockdown of *NOX4* using siRNA (Fig. S2B) rescued ferroptosis after erastin treatment in these cells (Figs. 2F and G). The high ROS levels in TD cells were also markedly reduced following NOX4 depletion (Fig. 2H). Similar results were obtained from the stable knockdown of NOX4 using shRNA (clone #4, Fig. S2C) (Figs. 2I and J).

### NOX4 as an insufficient marker for erastin sensitivity

Because the high-level expression of *NOX4* in TD cells appeared to be responsible for erastin-induced ferroptosis (Fig. 2), we hypothesized that cancer cells with high-level *NOX4* expression would be susceptible to erastin-induced ferroptosis. Given that a few previous studies have demonstrated the importance of *NOX4* in cancer malignancy [41], metastasis [42] and drug resistance [43] as well as in the regulation of EMT [44, 45], erastin could be a promising candidate drug for the treatment of *NOX4-*expressing cancers with an otherwise poor prognosis [41].

Because strong mitogenic signaling from RAS oncogenes produces ROS [46], which promote cell proliferation, it is possible that cancer cells with an RAS oncogene may be more susceptible to erastin [1, 2], leading to the suppression of GSH synthesis [47]. Thus, we examined erastin sensitivity in seven in-house lung cancer cell lines with and without RAS mutations (Fig. S3A). Unexpectedly, it was found that the oncogenic mutation of *KRAS* was not associated with either erastin sensitivity or basal ROS levels (Figs. 3A and S3B). The seven lung cancer cell lines could be classified into erastin-sensitive (erastin S: H1650, Calu1 and TD) and erastin-resistant (erastin R: H460, H1299, H358 and A549) groups regardless of the presence of a *KRAS* mutation or ROS levels (Figs. S3A and B). The time-dependent cell viability of Calu1 cells (the most sensitive to erastin of the tested cell lines) after erastin treatment was markedly restored by BME treatment and significantly delayed by ferrostatin treatment, confirming that Calu1 cells underwent erastin-induced ferroptosis (Fig. S3C). Similarly, pretreatment with GKT-137831 markedly rescued cell death in Calu1 cells (Fig. S3D), while the ectopic expression of *NOX4* in erastin-resistant A549 and H358 cells sensitized them to erastin treatment (Fig. S3E). These results are consistent with a recent study that demonstrated that the NOX4 induced by TAZ treatment leads to ferroptotic cell death [48].

To our surprise, the expression levels of *NOX4* and other *NOXs* (e.g. NOX1, 2, 3 and 5) were not strongly associated with erastin sensitivity (Fig. 3B), in contrast to a previous study that reported a close correlation between *NOX1* or *NOX4* expression and the response to erastin [3]. In addition, the expression of *ZEB1* and *GPX4* in these cell lines was not closely correlated with erastin sensitivity (Fig. 3C). This was also observed in lung cancer cell line data obtained from the CCLE and CTRP (Fig. 3D). As a result, we concluded that the individual expression levels of these genes cannot be used to indicate the susceptibility of cancer cells to erastin-induced ferroptosis.

### Systematic investigation of the molecular mechanisms associated with erastin response

To further examine the association between erastin sensitivity and known genomic characteristics in a variety of cancer cell lines, we explored cell-line omics profiles and drug response data from the CCLE and CTRP. Consistent with previous observations (Fig. 1), the expression profiles for the isogenic lung cancer cell models A549 and TD were clustered with those for the erastin R and S groups, respectively, of CCLE lung cancer cells (Fig. 4A, Fig. S4A). However, though erastin was initially employed to target oncogenic RAS, no consistent association between erastin sensitivity and status of RAS mutations was observed (Fig. S4B). Rather HRAS mutant cell lines exhibited moderate resistance to erastin (t-test, *P* < 0.05). In addition, a previous report has shown that therapy-resistant mesenchymal cancer cells that are highly sensitive to GPX4 inhibitors are also sensitive to erastin [9]. However, our correlation analysis between cell-line mesenchymal scores [9] and drug sensitivity found erastin had only a modest effect on the mesenchymal cancer cells (Fig. S4C). In fact, the sensitivity to erastin differed from the sensitivity to GPX inhibitors according to the cell type is still in question [49]. Taken together, neither RAS mutations nor mesenchymal signatures are suitable as indicators of erastin sensitivity.

We subsequently employed a two-step process to determine the molecular pathways that contribute to erastin sensitivity using pan-cancer transcriptome data. First, we conducted single-sample gene set enrichment analysis (ssGSEA) across 598 non-hematologic cancer cell lines using biological pathway information, which yielded 443 pathway-enrichment scores (PESs) for each cell line (Table S1). By correlating these cell-line PESs with sensitivity to erastin, the 16 pathways most strongly related to erastin sensitivity were selected (Fig. 4B). The A549 and TD cell lines exhibited similar enrichment patterns for these pathways (Fig. 4C). In the second step, we applied elastic net regularized regression [31] to identify the subset of genes that were most strongly associated with the response to erastin in each of the 16 pathways. Overall, regularized regression outperformed general linear regression and ssGSEA (Fig. 4D, Table S2), suggesting that erastin sensitivity can be predicted by the integrated expression of a set of designated genes in these pathways. In particular, the elastic net based on the nuclear receptor meta-pathway (hereafter referred to as the NRM) had the strongest correlation (*r* = 0.456), which was surprisingly higher than that based on all genes (*r* = 0.429, the red line in Fig. 4D).

### The relationship between nuclear receptors and erastin resistance as revealed by predictive models

To identify the most predictive gene signature in the NRM, we constructed an elastic net model using all 598 cancer cell lines employing the expression of the 312 genes involved in the NRM. This model included 43 predictor genes with non-zero coefficients (Fig. 5A), most of which were involved in one or more of the other 15 enriched pathways (Fig. 5B). Among them, NRF2 and Aryl hydrocarbon receptor (AhR) pathways accounted for the largest proportion (Fig. 5C and Fig. S5A). Correlation analysis of the cell-line PESs and sensitivity to class II FINs (e.g. RSL3, ML162, and ML210, which are direct *GPX4* inhibitors) revealed that NRF2- and oxidation-related genes were commonly associated with resistance to FINs (Fig. S5B). This is in line with previous reports that found that NRF2, a master regulator of oxidative stress responses, modulates ferroptosis [50, 51]. However, the AhR pathway was found to be the only factor associated with resistance to erastin, unlike the NRF2 pathway, which is commonly associated with resistance to both erastin and class II FINs (Fig. 5D). Interestingly, an analysis of genome-wide CRISPR-Cas9 loss-of-function screening data (DepMap 19Q2, https://depmap.org/portal/) revealed the high dependency on GPX4 or genes encoding selenoproteins (*SEPSECS, EEFSEC*, and *SEPHS2*) in cells that are sensitive to all FINs (Fig. 5E, Fig. S5C), consistent with previous studies [10, 52]. In contrast, the knockout of *AHR* (the gene encoding AhR) led to vulnerability in erastin-resistant cancer cell lines, while the deficiency of *NFE2L2* (the gene encoding NRF2) increased sensitivity to both erastin- and GPX4 inhibitors-resistant cells (Fig. S5C). This suggests that the AhR signature, which complements the NRF2 signature, accounts for the unique dependency of erastin-resistant cells, thus increasing the predictive power of the NRM model for erastin response.

**Figure 5.**
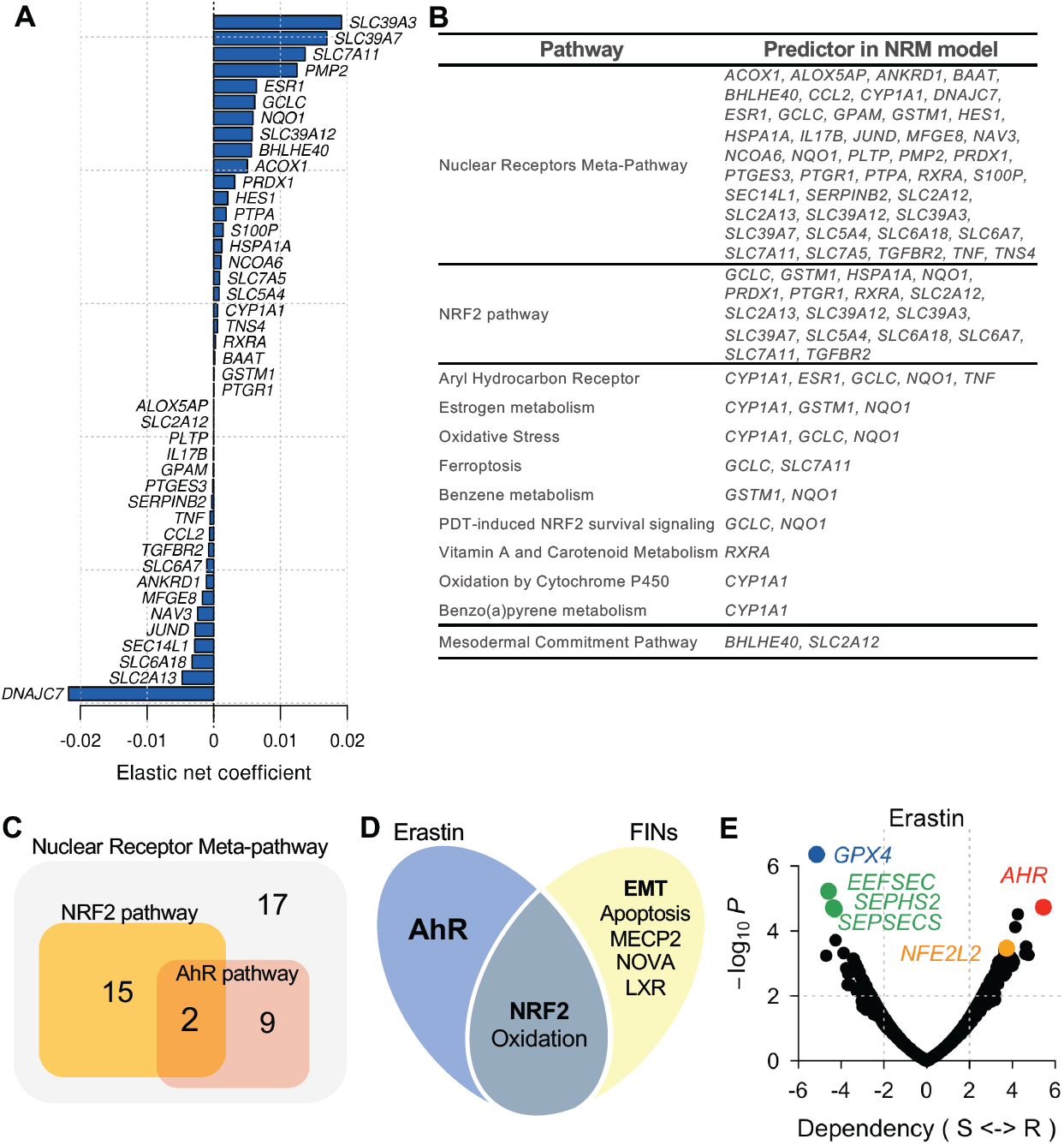
Association of nuclear receptors with erastin resistance as revealed by the predictive models. **(A)** Bar plot showing the weight of the 43 predictor genes in the model for erastin sensitivity. **(B)** NRM model predictors involved in each of the top pathways. **(C)** Venn diagram of the number of shared genes involved in the NRF2 and AhR pathways from among the 43 predictor genes in the NRM model. **(D)** Venn diagram of representative functional terms accounting for the response to erastin and FINs (RSL3, ML210, and ML162). **(E)** Volcano plot highlighting CRISPR hits associated with erastin sensitivity across pan-cancer cell lines. Dependency is defined as the *t*-statistic calculated by testing the difference between erastin sensitivity (AUC) in the non-dependent and dependent cell lines for the corresponding gene.

### High correlation between NRF2 activity and erastin resistance

As predicted, the basal NRF2-dependent gene response determined by the antioxidant response element (ARE) was significantly stronger in erastin R cancer cells than in erastin S cancer cells (Fig. 6A). In the isogenic pair with different sensitivity to erastin (A549 vs TD cells), induction of *CHAC1*, an NRF2 downstream gene [53] that acts as a marker for ferroptosis [54], was significantly lower in TD cells (Fig. S6A). In addition, the activation of the NRF2-dependent gene response by tert-Butylhydroquinone (tBHQ), a well-established NRF2 activator [55], induced typical NRF2-dependent genes such as *GCLC, GCLM*, and *NQO1* in a dose-dependent manner (Fig. 6B). Under these conditions, the erastin sensitivity of Calu1 cells was significantly reduced (Fig. 6C). Conversely, the knockdown of *NFE2L2* (which encodes NRF2) in erastin R cells sensitized them to erastin treatment (Fig. 6D). These results suggest that the NRF2 pathway is closely associated with erastin sensitivity. Similarly, the mutation of *KEAP1*, which leads to an NRF2-dependent adaptive response in cancers [56] was significantly correlated with erastin resistance, while *RAS, TP53*, or *NFE2L2* mutations were not (Fig. S6B). The expression of six typical NRF2 target genes (*SLC7A11*, a molecular target of erastin, *NQO1, GCLC, GCLM, ME1*, and *SRXN1*) involved in anti-oxidant activity was significantly correlated with erastin sensitivity in lung cancer cells and all cancer cells (Fig. S6C). These results support that conjecture that NRF2-associated genes are a major determinant of erastin sensitivity.

**Figure 6.**
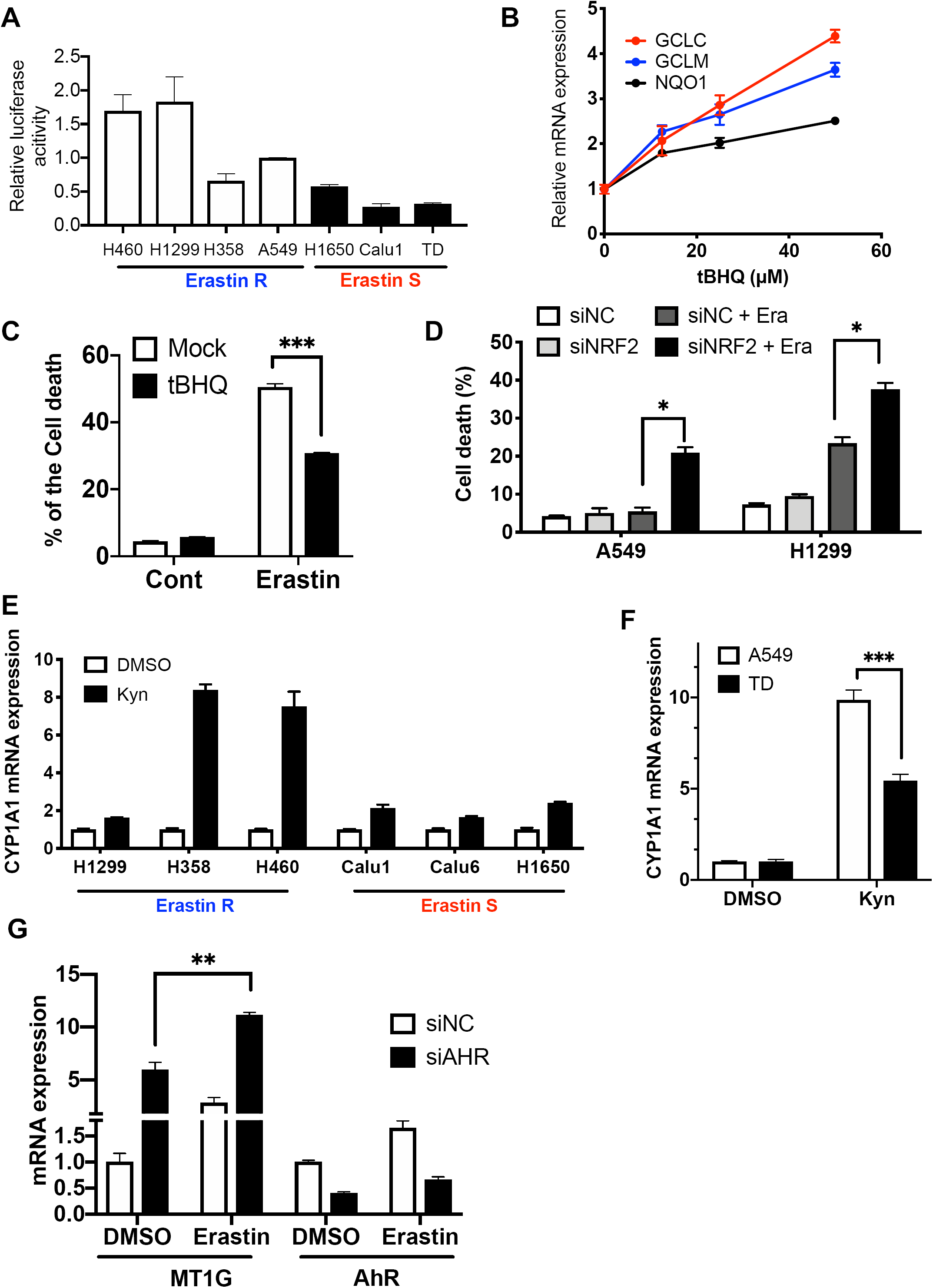
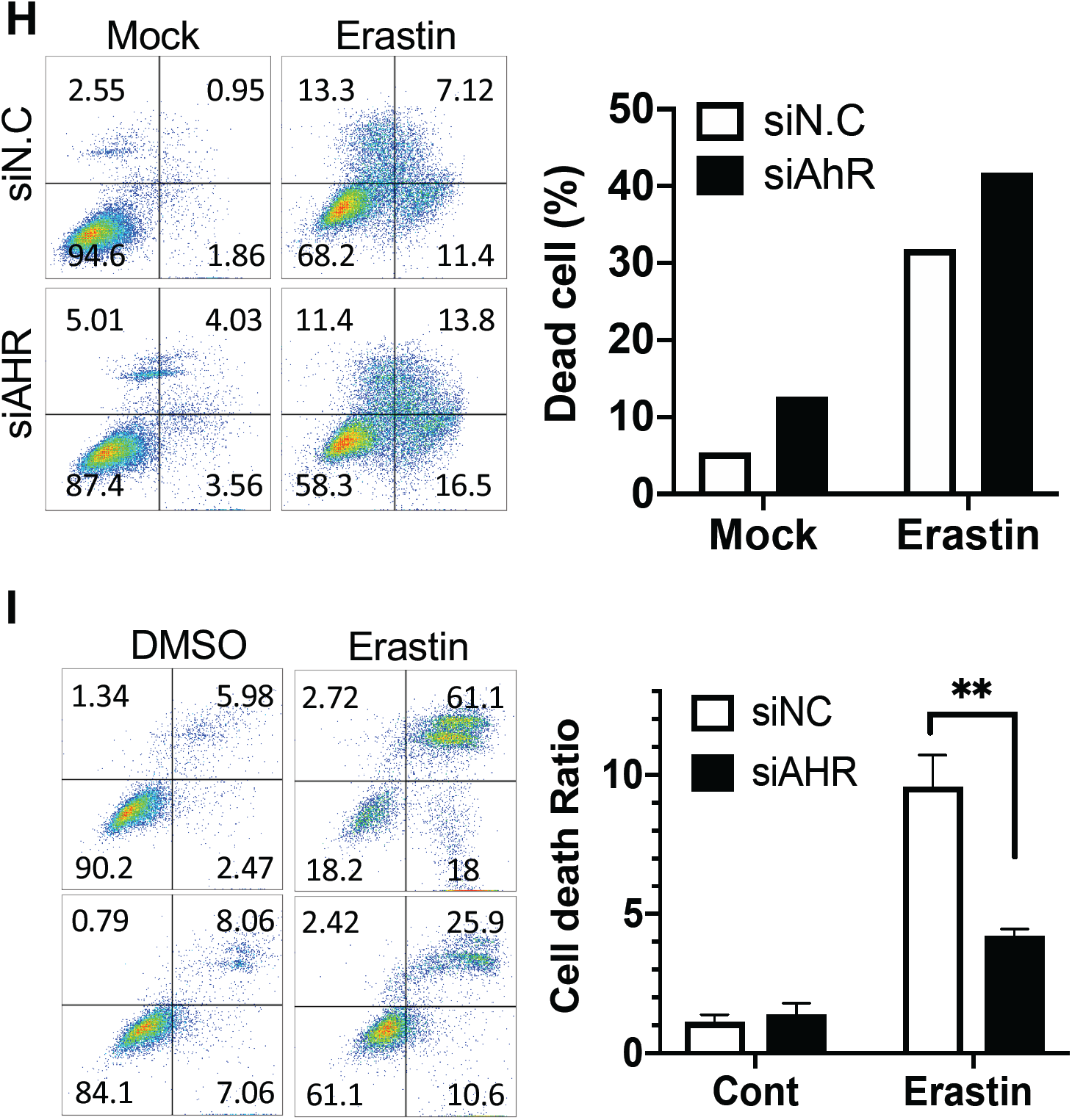
High correlation between NRF2 activity and erastin resistance. **(A)** Relative luciferase activity of the NRF2 promoter in the indicated lung cancer cell lines. **(B)** Relative mRNA levels of NRF2 downstream genes (*GCLC, GCLM*, and *NQO1*) 24 hours after tBHQ treatment at the indicated concentrations in Calu1. **(C)** Graphical representation of cell death population size 24 hours after erastin treatment with 50 μM tBHQ treatment. **(D)** Cell death population size 24 hours after erastin treatment in the control (siNC) or NRF2-knockdown cells with siRNA (siNRF2) in A549 and H1299 cell lines. **(E, F)** mRNA expression of *CYP1A1* in the indicated lung cancer cell lines (E) and the isogenic pairing of A549 and TD cells (F) with or without kynurenine (Kyn: 100 nM). **(G)** Relative mRNA expression of *MT1G* in A549 8 hours after 80 μM erastin treatment in *AHR*-knockdown cells with siRNA transfection. **(H, I)** Cell death population with Annexin V positive cells 24 hours after 200 μM erastin treatment.

### Association between AhR dependency and erastin resistance

Because the AhR signature was enriched in the NRM model, we sought to determine whether AhR activity differed between the erastin R and S groups. To achieve this, AhR activity was monitored in three cell lines each within the erastin R and S groups by measuring *CYP1A1*, a well-characterized AhR downstream target, after treatment with kynurenine (Kyn), a ligand of AhR [57]. As predicted, *CYP1A1* was strongly induced by Kyn treatment in two out of the three cell lines in the erastin R group, while it was only moderately induced in all three cell lines in the erastin S group (Fig. 6E).

Modulation of the AhR gene response was then compared for the isogenic pair of A549 and TD cells with different erastin sensitivities. Consistent with the data shown in Figure 6E, the AhR gene response following Kyn treatment was markedly lower in TD cells, which showed higher sensitivity to erastin, compared to that of A549 (Fig. 6F). Similar results with AhR activation with Kyn (Fig. S6D) and high dose of tBHQ [58] (Fig. S6F) were reproduced in the comparison between A549 and Calu1. These observations led us to hypothesize that AhR activation in A549 may confer erastin resistance. Based on the increase in *MT1G*, which is induced during ferroptosis [59] (Fig. 6G), and in the number of dead cells (Fig. 6H) following AhR depletion and subsequent erastin treatment, we concluded that the depletion of AhR promoted ferroptotic cell death in A549 cells. In contrast, erastin sensitivity was attenuated by AhR depletion in Calu1 (Fig. 6I). Thus, the enrichment of the AhR pathways in the NRM model may account for the clear dependency of erastin sensitivity on AhR.

### Predictive performance of the NRM model for erastin sensitivity

Next, we assessed the specificity of the NRM signature in terms of predicting erastin response. Correlating cell-line NRM predictions with the sensitivity to each of 543 drugs in CTRP showed that erastin, followed by class II FINs (ML210, RSL3, ML162, and ML239), had the highest priority, but STATINs that have been shown to induce ferroptosis [9] did not (Fig. 7A). The NRM gene signature was more effective in predicting erastin sensitivity than the three independent mesenchymal signatures used to identify FINs as the best treatment option for mesenchymal cancer in a previous study [9] (*P* = 7.13 × 10^−7^, Fig. 7B). These results together suggest that the NRM signature allows specific predictions of cellular vulnerability to erastin-induced ferroptosis to be made, but not to FIN- or STATIN-induced ferroptosis.

**Figure 7.**
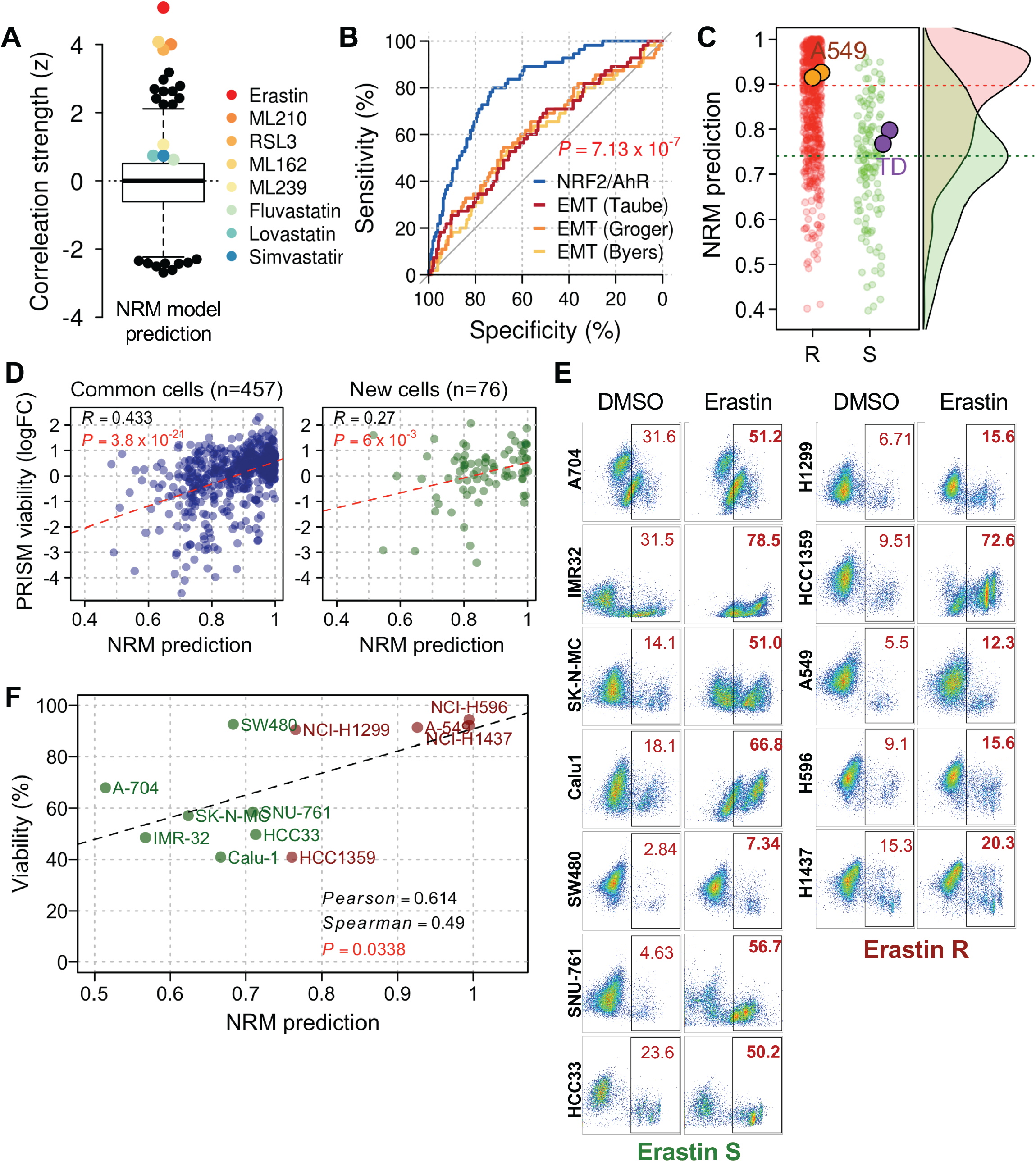
Evaluation of the NRM model for erastin sensitivity. **(A)** Prioritization of compounds based on the correlation between NRM-based LOOCV predictions and sensitivity profiles (AUC) for 543 compounds in the CTRP. **(B)** Receiver operating characteristic (ROC) curve illustrating the performance of the erastin sensitivity predictions using the NRM-based model (ROCAUC=0.84) and three mesenchymal scores obtained from a previous study. A cell with an AUC lower than 0.7 was considered to be sensitive. **(C)** Distribution of NRM prediction scores for all CCLE cancer cell lines, A549, and TD cells. Cell-line R and S groups were determined by the AUC measured for each cell line (higher or lower than 0.7, respectively). **(D)** Comparison of NRM predictions to PRISM cell viability data (log2 fold change) with 2.5 μM erastin treatment. Primary screened data were obtained from https://depmap.org/repurposing. **(E)** Flow cytometry plot for the 7-AAD positive cell population size 24 hours after 120 μM erastin treatment in the indicated cell lines. **(F)** Comparison of NRM predictions to cell viability with 7-AAD positive cells after 120 μM erastin treatment in the indicated cell lines.

To evaluate the predictive performance using independent datasets, we applied the NRM model to gene expression data for A549 and TD and showed accurate predictions of erastin sensitivity in each cell line (Fig. 7C). We then compared our NRM prediction to recently published drug response data obtained from PRISM viability assay, a multiplexing screening with molecular barcoding method [26, 60]. The PRISM dataset provides an erastin-sensitivity profile across 533 cancer cell lines, 457 of which have also been screened in the CTRP. Encouragingly, the PRISM profile was more closely correlated with the NRM prediction (Spearman’s r = 0.433) than the CTRP profile (Spearman’s r = 0.358) (Fig. 7D, left panel). We also observed overall agreement across the 76 cancer cell lines present only in the PRISM dataset (Spearman’s r = 0.27, *P*=0.006) (Fig. 7D, right panel). For further validation, we additionally predicted the erastin sensitivity of 334 cancer cell lines not used in NRM modeling and selected nine test cell lines and three control cell lines (Fig. S7A, Table S3). The nine test cell lines were divided into responders (erastin S) and non-responders (erastin R) based on the NRM prediction of the control cell lines. The cell death population size of twelve cancer cell lines was examined after erastin treatment (Fig. 7E). The erastin sensitivity of seven of the cancer cell lines (all except SW480 and HCC1359) was highly correlated with the NRM prediction (Fig. 7F). These results provide evidence that our NRM model is universally applicable, being able to accurately predict erastin sensitivity based on transcriptome data generated from different cohorts.

## Discussion

There is emerging evidence that mesenchymal-type cancer cells are responsible for malignant phenotypes [61, 62]. Thus, EMT-associated molecular targets that govern chemoresistance [27, 37, 63] or metastatic potential [35, 36] have been extensively studied for the development of novel anti-cancer therapies [64]. As such, the induction of the selective death of mesenchymal-type cancer cells using small molecules identified via library screening [65] or *in silico* gene signature-based analysis [9] has been highlighted as an effective potential strategy.

Using isogenic lung cancer cell models, we observed that selective ferroptosis occurred in chemoresistant mesenchymal lung cancer cells [27, 37] (Fig. S1A) following erastin treatment (Fig. 1) due to the redox imbalance caused by the high expression of *NOX4* and subsequent depletion of GSH (Fig. 2). However, the mRNA expression levels of both *NOX4* and previously identified determinants (e.g. *GPX4, ZEB1*, and other *NOXs*) were unable to be used as indicators of erastin sensitivity in other lung cancer cells due to cell-to-cell variation (Fig. 3). Therefore, we adopted a pharmacogenomic approach that utilized pan-cancer cell-line omics to fully explore predictive biomarkers for erastin sensitivity. We initially examined whether known markers such as the *KRAS* mutation status or mesenchymal signatures explained erastin sensitivity, but neither were able to predict erastin sensitivity in either lung cancer or pan-cancer cell lines.

In our study, we applied a two-step strategy in which the molecular pathways associated with the erastin response were screened and then the 16 top pathways were assessed to identify the most relevant biomarkers. Interestingly, we found that the NRF2 and AhR pathways were strongly associated with erastin resistance in pan-cancer cell lines (Figs 4 and 5). The high dependency on the NRF2 pathway for the conferral of erastin sensitivity was then biochemically proven in cancer cell line models (Figs. 6). In particular, a diverse range of cancer malignancies results from continuous ROS generation [66] following oncogenic *RAS* mutations or elevated MAPK signaling [46], while excessive ROS can be sensitized to chemotherapeutics [67]. Thus, cancer cells may adapt to a high-ROS environment through the induction of anti-oxidant mechanisms, including KEAP1-NRF2 [68, 69], which is why NRF2 has been studied as a promising molecular target within advanced cancers. Moreover, the AhR gene response, which was induced by erastin treatment, was higher in erastin R cells than erastin S cells. Furthermore, the depletion of AhR sensitized only erastin R cells to erastin, while it desensitized erastin S cells (Fig. 6).

The NRM model, with the enriched NRF2 and AhR signatures used as predictors, has the potential to readily predict the erastin response of any cell lines whose transcriptome data are available. The robustness of this model was assessed using an independent pharmacogenomic dataset for pan-cancer cells and in-house isogenic lung cancer cells. The model was further experimentally validated using nine additional cancer cell lines whose erastin responsiveness had not yet been determined (Fig. 7). Given that an erastin analog is currently undergoing clinical trials for anti-cancer therapy [22], this approach would be useful for patient stratification in these trials to maximize their efficacy and for the selection of those patients most likely to respond to erastin-based anti-cancer therapy in the future.

## Data availability

RNA-seq of A549 and TD cell lines can be obtained from Gene Expression Omnibus (GEO) under accession number GSE135402.

## Code availability

Source code for training and testing the elastic net regression model is available from the corresponding authors upon request.

## Authors’ contributions

HJ.C conceived the overall study design, led the experiments and wrote the manuscript. H.L conceived the overall study design, performed bioinformatics analysis and wrote the manuscript. OS.K conducted the experiments, data analysis and wrote the first draft. EJ.K, HJ.K, JY.C, and YJ.K validated the experiments and generated cell lines. W.K discussed and interpreted the data. All authors contributed to manuscript writing and revising, and endorsed the final version of this manuscript.

