## Supplementary figures and figure legends for "Systematic identification of a nuclear receptor-enriched predictive signature for erastin-induced ferroptosis"

**Running title: Predictive gene signature for erastin sensitivity**

This PDF file includes:

Supplementary Materials and Methods

Supplementary Figures 1-6

**Supplementary Materials and Methods**

**Reagents**

Erastin (HY-15763) was purchased from MedchemExpress. Beta-mercaptoethanol (21985-023) was purchased from Thermo Fisher Scientific. N-acetylcysteine (A7250), GSH-MEE (G1404) was purchased from Sigma-Aldrich. Ferrostatin-1 (sc-498126), Z-VAD (sc-3067) was purchased from Santa Cruz Biotechnology Inc. shRNA targeting *NOX4* (SHCLANG-NM_016931) were purchased from Sigma Aldrich. Primary antibodies against cleaved caspase 3 (9664S) and β-actin (sc-47778) were obtained from Cell Signaling Technology and Santa Cruz, respectively.

**Annexin-V & 7-AAD staining**

Annexin-V & 7-AAD staining to determine the population of dead cells was performed according to the manufacturer's instructions (559763, BD Pharmingen). The stained cells were subjected to flow cytometry and the results were analyzed using a Becton-Dickinson FACS Calibur-1 with the Cell-QUEST software.

**DCF-DA staining**

2',7'-dichlorofluorescin diacetate (DCF-DA, D6883, Sigma-Aldrich) staining to determine the Cellular ROS level was performed according to the manufacturer's instructions. The stained cells were subjected to flow cytometry and the results were analyzed using a Becton-Dickinson FACS Calibur-1 with the Cell-QUEST software.

**GSH fluorometric assay**

Cells were seeded about 1 x $10^3$ in 96-well white plate. Detection of total GSH and reduced/oxidized GSH ratio was performed according to the manufacturer's instructions (VV6611, promega).

**Live detection of GSH/GSSG ratio by fluorescent probe**

Cells were seeded about 1 x $10^3$ in 96-well black plate. FreSHtracer (Cell2in), fluorescent probe that combines with GSH to show 510nm wavelength and with GSSG to show 580nM wavelength, was treated to plate. Intensity of fluorescence from cells was detected by Opperatta High-Content Imaging system (HH12000000, PerkinElmer).

**Cell culture**

A549, TD and H1299 cell lines were maintained in Dulbecco's modified Eagle's medium (DMEM). H358, H460 cell lines were maintained in Roswell Park Memorial Institute medium (RPMI) 1640. Calu1 cell line was maintained in McCoy's 5A medium. DMEM, RPMI and McCoy's media were supplemented with 10% (v/v) fetal bovine serum (FBS), gentamicin (50 μg/ml) at 37°**C** in a humidified atmosphere of 5% $CO_2$ in the air.

**Total RNA extraction and Quantitative real time PCR**

Total RNA was extracted by Total RNA Extraction Kit (17061, Intron). Total RNA was converted to cDNA using RT Master Mix (RR036, TAKARA) based on the manufacturer's instruction. The synthesized cDNAs were used as template to examine the real-time PCR using Light Cycler 480 Instrument II (Roche) with TB Green (RR420) under TAKARA's 2 step protocol.

**Transfection**

Transient transfection was performed with Lipofectamine 2000 (#11668019, Invitrogen) according to the manufacture instruction. siRNA targeting NOX4, AHR and NFE2L as well as the non-targeting control siRNA were purchased from Bioneer. siRNA transfection was performed using DharmaFECT (#T-2001-02, Dharmacon Inc.) in accordance to the manufacture instruction.

**Dual luciferase assay**

Cells were transfected with specific promoter-luciferase vector and pRL vector using above description. Cells lysate was extracted with 1X lysis buffer (5X lysis buffer diluted in ultrapure water to 1X) for 20 minutes. Cell pellet were downed through 13,000 RPM for 20 minutes, and supernatant was transferred to E-tube. Then, reporter assay was performed according to the Dual-Luciferase Reporter Assay System (E1980, Promega).

**Supplementary Figure legends**

**Figure S1 (A)** List of genesets, significantly enriched in TD compared to A549 cells. **(B)** A graphical scheme (upper) and merged image of GFP-tagged A549 and TD at indicated timepoint after erastin treatment.

**Figure S2 (A)** Volcano plot showing gene expression comparison between TD and A549 cells. ROS-related genes are indicated in red (upregulated) or green (downregulated). **(B)** Relative mRNA expression level of *NOX4* in A549 and TD cell with *NOX4* siRNA transfection. **(C)** Relative mRNA levels of *NOX4* after knockdown with each different shRNA, was shown in a bar graph.

**Figure S3 (A)** List of in house lung cancer cell lines with erastin sensitivity and *KRAS* mutation status. **(B)** Histogram plot of flow cytometry for DCF-DA of each cell lines (Blue: erastin resistant, red: erastin sensitive, * for *KRAS* mutation). **(C)** Cell viability rate of TD cell with indicated treatment was graphically represented (F: ferrostatin). **(D)** Graphical presentation of cell death 24 hours after erastin (40μM) and GKT-137831 (20μM) in Calu1 and H460 cell. **(E)** Percentage of cell death population with 40uM of erastin treatment after induction of *NOX4* overexpressing vector in A549 and H358.

**Figure S4 (A)** CCLE cancer cell lines ranked by erastin sensitivity. Distribution of erastin sensitivity (AUC) is shown in the right panel. Four erastin resistant cells (NCI-H1299, A549, NCI-H359, and NCI-H460) and two erastin sensitive cells (Calu1 and NCI-H1650) are indicated in green and red, respectively. **(B)** Distribution of erastin sensitivity (AUC) according to mutation status (Wild type: Wt, and Mutant: Mut) of *KRAS* and *HRAS* in 598 non-hematological cancer cell line. The number of cell lines corresponding to each status is denoted below. *P*-values were calculated with the two-tailed t-tests. **(C)** Box plot showing the extent of correlation between cell-line mesenchymal scores and AUCs for each compound in CTRP. Correlation strength is defined as the *z* scored Pearson's correlation coefficient as described previously [1].

**Figure S5 (A)** Venn diagram showing genes involved in the NRF2 and AhR pathways from among the 43 predictor genes in the NRM model. **(B)** The pathways most closely associated with the response to erastin and FINs (RSL3, ML210, and ML162). The association between the pathways and drug sensitivity was measured using Pearson's correlation between the cell-line PESs and AUC. Pathways with an absolute *z*-normalized correlation coefficient greater than 2 were selected. Positive and negative correlations are shown in red and green, respectively. Gene annotations for the 445 pathways were obtained from Wikipathways. Representative functional terms accounting for cell-line responses to erastin, RSL3, ML210, and ML162, respectively, are indicated by yellow boxes. **(C)** Volcano plot highlighting CRISPR hits associated with the response to erastin and FINs (RSL3, ML210, and ML162) across pan-cancer cell lines. Dependency is defined as the *t*-statistic calculated by testing the difference between erastin sensitivity (AUC) in the non-dependent and dependent cell lines for the corresponding gene.

**Figure S6 (A)** mRNA expression of *MT1G* 24 hours after erastin treatment in A549 and TD cell. **(B)** Distribution of erastin sensitivity (AUC) according to mutation status (Wild type: Wt, and Mutant: Mut) of indicative genes in 598 non-hematological and 124 lung cancer cell line. The number of cell lines corresponding to each status is denoted below. *P*-values for significance from two-tailed t-tests. **(C)** Relationship between erastin sensitivity (AUC) and gene expression values (logTPM) of six typical NRF2 downstream target genes. Lung cancer cell lines are denoted in red, and others in black. Erastin sensitivity data from CTRP and gene expression from CCLE RNA-seq ($log_2$TPM) is shown. **(D)** Relative mRNA expression of *CYP1A1* 24 hours after indicative dose of tBHQ treatment in A549 and Calu1. **(E)** *CYP1A1* expression after 3 hours of kynureine (Kyn:100nM) treatment after *AHR* knockdown by siRNA treatment in A549 and Calu1.

**Figure S7 (A)** List of twelve cancer cell lines with NRM prediction score (S.P.C: positive control for erastin sensitive cell line, R.P.C: negative control for erastin resistant cell line).

### Figure S1

## A

| GSEA | ES | NES | FDR | Data origin |
|---|---|---|---|---|
| **EMT-related gene signatures** | | | | |
| EMT | 0.64 | 2.35 | < 0.001 | Hallmark |
| EMT | 0.54 | 1.79 | 0.009 | GO BP |
| EMT (Up-regulated genes) | 0.64 | 2.17 | < 0.001 | ANASTASSIOU |
| EMT (Up-regulated genes) | 0.47 | 1.61 | 0.014 | GOTZMANN |
| EMT (Up-regulated genes) | 0.46 | 1.57 | 0.019 | JECHLINGER |
| Mesenchymal cell differentiation | 0.46 | 1.66 | 0.022 | GO BP |
| Mesenchymal cell proliferation | 0.51 | 1.56 | 0.043 | GO BP |
| Mesenchyme development | 0.43 | 1.59 | 0.035 | GO BP |
| Mesenchyme morphogenesis | 0.52 | 1.68 | 0.02 | GO BP |
| Angiogenesis | 0.54 | 1.71 | 0.001 | Hallmark |
| **Therapy resistance-related gene signatures** | | | | |
| Dasatinib resistance (Up genes) | 0.60 | 2.09 | < 0.001 | Huang et al. |
| Tamoxifen resistance (Up genes) | 0.63 | 1.79 | 0.002 | Masri et al. |
| BCL2 inhibitor resistance (Up genes) | 0.57 | 1.74 | 0.004 | Hann et al. |
| Alkylating agents resistance (Up genes) | 0.51 | 1.52 | 0.027 | Bacolod et al. |
| Tamoxifen resistance (Up genes) | 0.44 | 1.5 | 0.033 | Riggins et al. |
| Cisplatin resistance (Up genes) | 0.49 | 1.48 | 0.039 | Li et al. |
| Doxorubicin resistance | 0.44 | 1.46 | 0.044 | Gyorffy et al. |
| Gamma radiation resistance | 0.52 | 1.48 | 0.038 | Amundson et al. |

| GSEA | ES | NES | FDR | Data origin |
|---|---|---|---|---|
| **Cancer signaling gene signatures** | | | | |
| Hedgehog signaling | 0.57 | 1.81 | < 0.001 | Hallmark |
| TGF β signaling | 0.52 | 1.71 | 0.001 | Hallmark |
| TNFα signaling via NF-κB | 0.42 | 1.57 | 0.003 | Hallmark |
| Wnt/β-catenin signaling | 0.47 | 1.50 | 0.005 | Hallmark |
| IL-6/JAK/STAT3 signaling | 0.38 | 1.34 | 0.028 | Hallmark |
| KRAS signaling (Up genes) | 0.36 | 1.33 | 0.027 | Hallmark |
| Notch signaling | 0.41 | 1.31 | 0.036 | Hallmark |
| NF-κB signaling | 0.63 | 2.00 | < 0.001 | Schoen |
| CD40 signaling (Up genes) | 0.48 | 1.68 | 0.007 | Basso et al. |
| IL6 signaling (Up genes) | 0.5 | 1.66 | 0.008 | Dasu et al. |
| TNF signaling (Up genes) | 0.44 | 1.53 | 0.027 | Sana et al. |
| BMP signaling | 0.53 | 1.82 | 0.007 | GO BP |
| Non canonical WNT signaling | 0.62 | 1.79 | 0.009 | GO BP |
| VEGFR signaling pathway | 0.57 | 1.72 | 0.015 | GO BP |
| GPCR signaling pathway | 0.62 | 1.72 | 0.015 | GO BP |
| Purinergic receptor signaling pathway | 0.58 | 1.67 | 0.02 | GO BP |
| Inositol phosphate mediated signaling | 0.58 | 1.67 | 0.02 | GO BP |
| Glutamate receptor signaling pathway | 0.51 | 1.63 | 0.028 | GO BP |
| Integrin mediated signaling pathway | 0.47 | 1.62 | 0.03 | GO BP |
| IGFR signaling pathway | 0.55 | 1.55 | 0.047 | GO BP |

## B

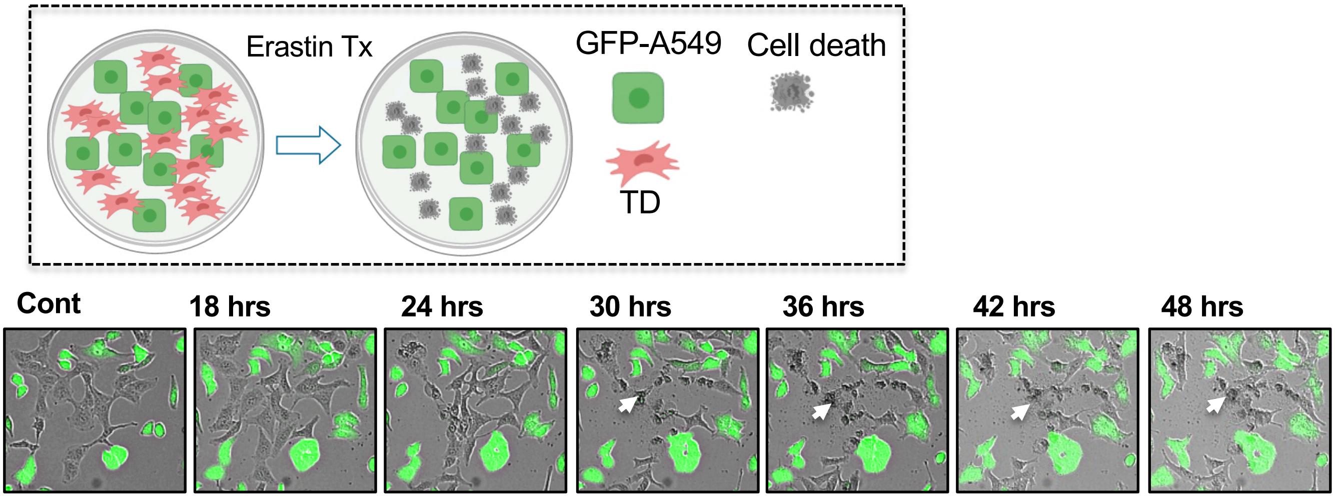

### Figure S2

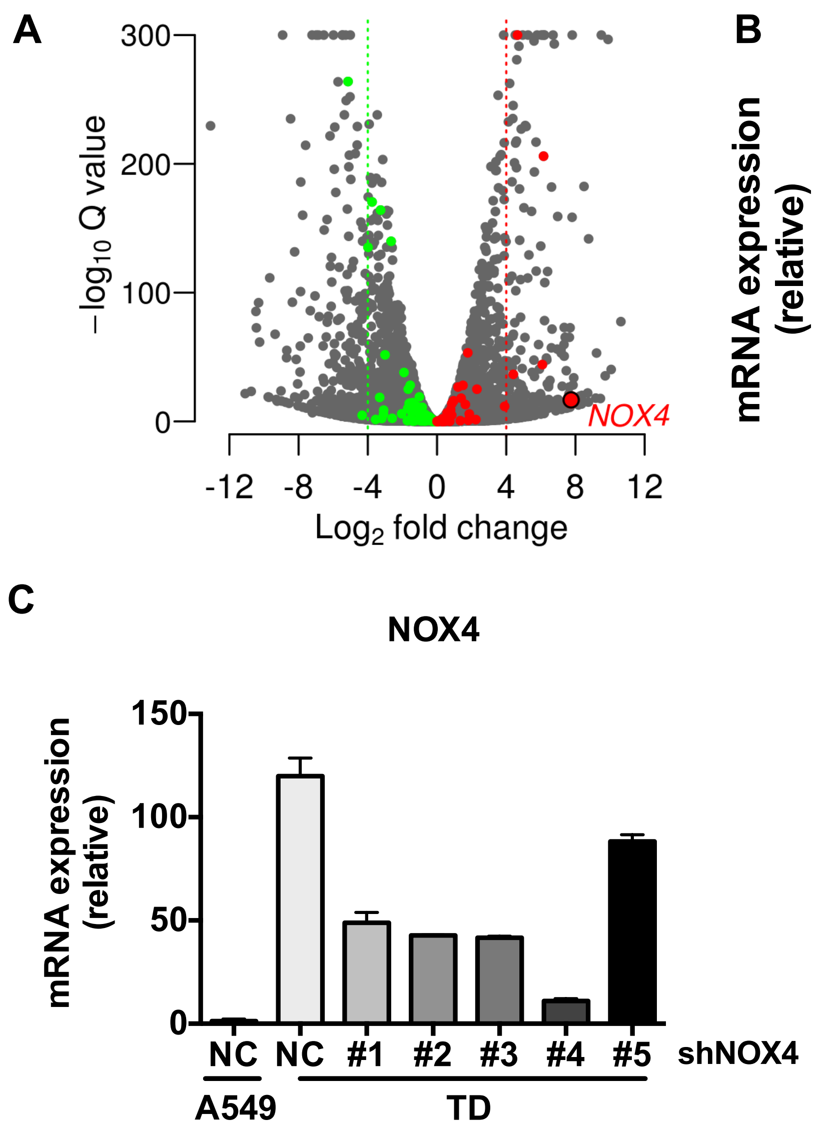
A
300
200
100
0
−log10 Q value
-12
-8
-4
0
4
8
12
Log2 fold change
NOX4
B
NOX4
1000
800
600
400
200
0
mRNA expression (relative)
siNC
siNOX4
****
A549
TD
C
NOX4
150
100
50
0
mRNA expression (relative)
NC
NC
#1
#2
#3
#4
#5
shNOX4
A549
TD

#### Figure S3

### A

| Cell lines | KRAS | Erastin Sensitivity |
|---|---|---|
| H460 | Missense | R |
| H1299 | Wt | R |
| H358 | Missense | R |
| A549 | Missense | R |
| H1650 | Wt | S |
| Calu1 | Missense | S |
| TD | Missense | S |

### B

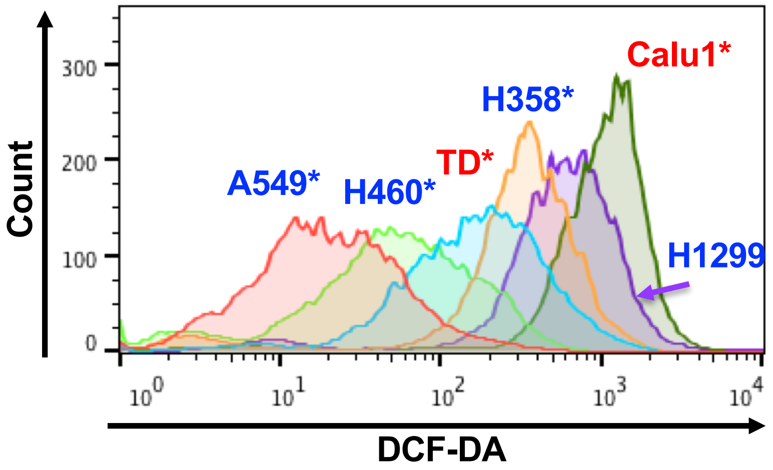


### C

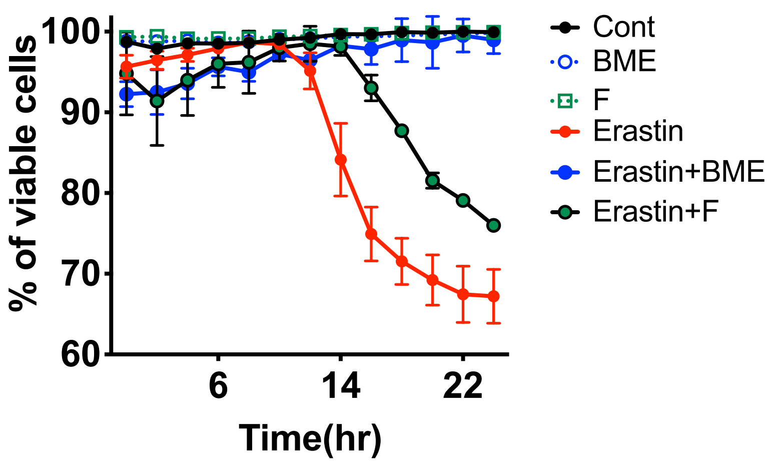


### D

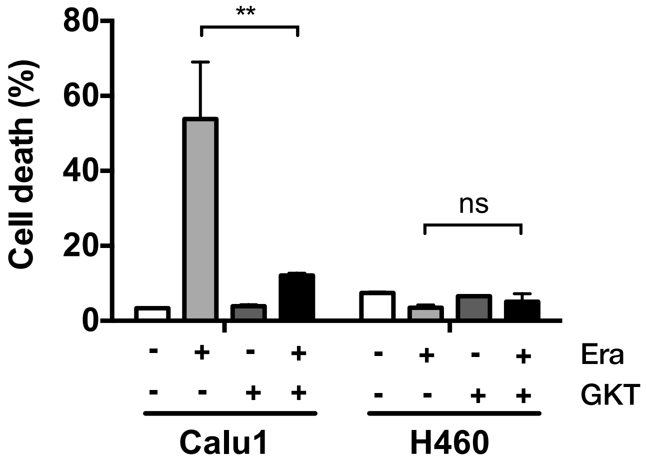


### E

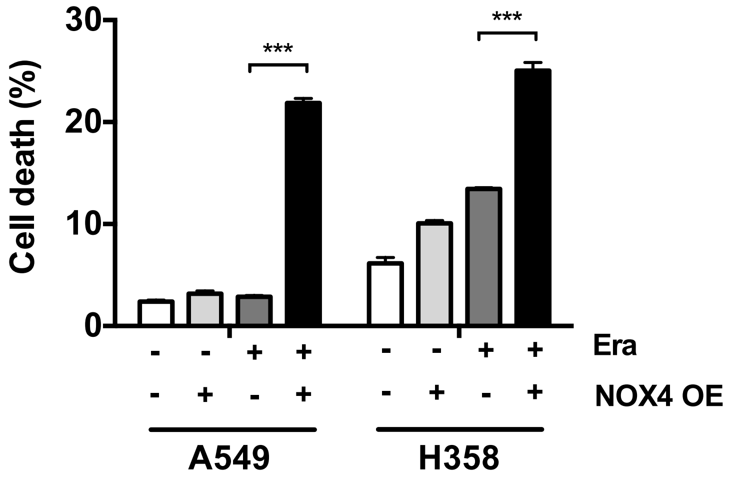

### Figure S4

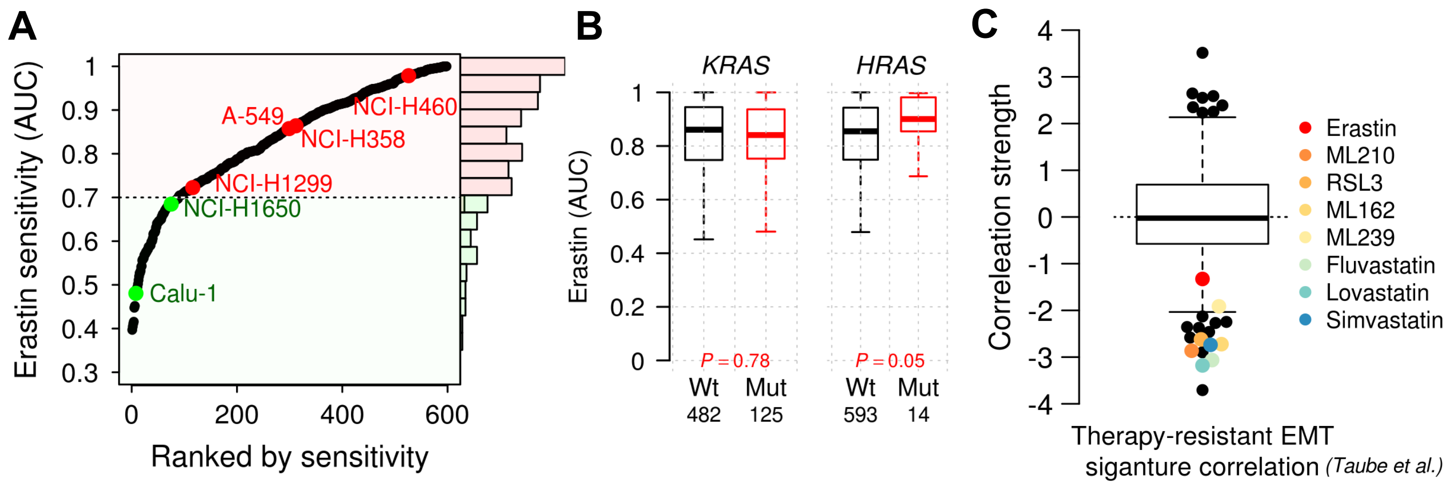
A
B
C
Erastin sensitivity (AUC)
Ranked by sensitivity
A-549
NCI-H460
NCI-H358
NCI-H1299
NCI-H1650
Calu-1
KRAS
HRAS
Erastin (AUC)
P = 0.78
P = 0.05
Wt
Mut
482
125
593
14
Correleation strength
Therapy-resistant EMT
siganture correlation (Taube et al.)
Erastin
ML210
RSL3
ML162
ML239
Fluvastatin
Lovastatin
Simvastatin

### Figure S5

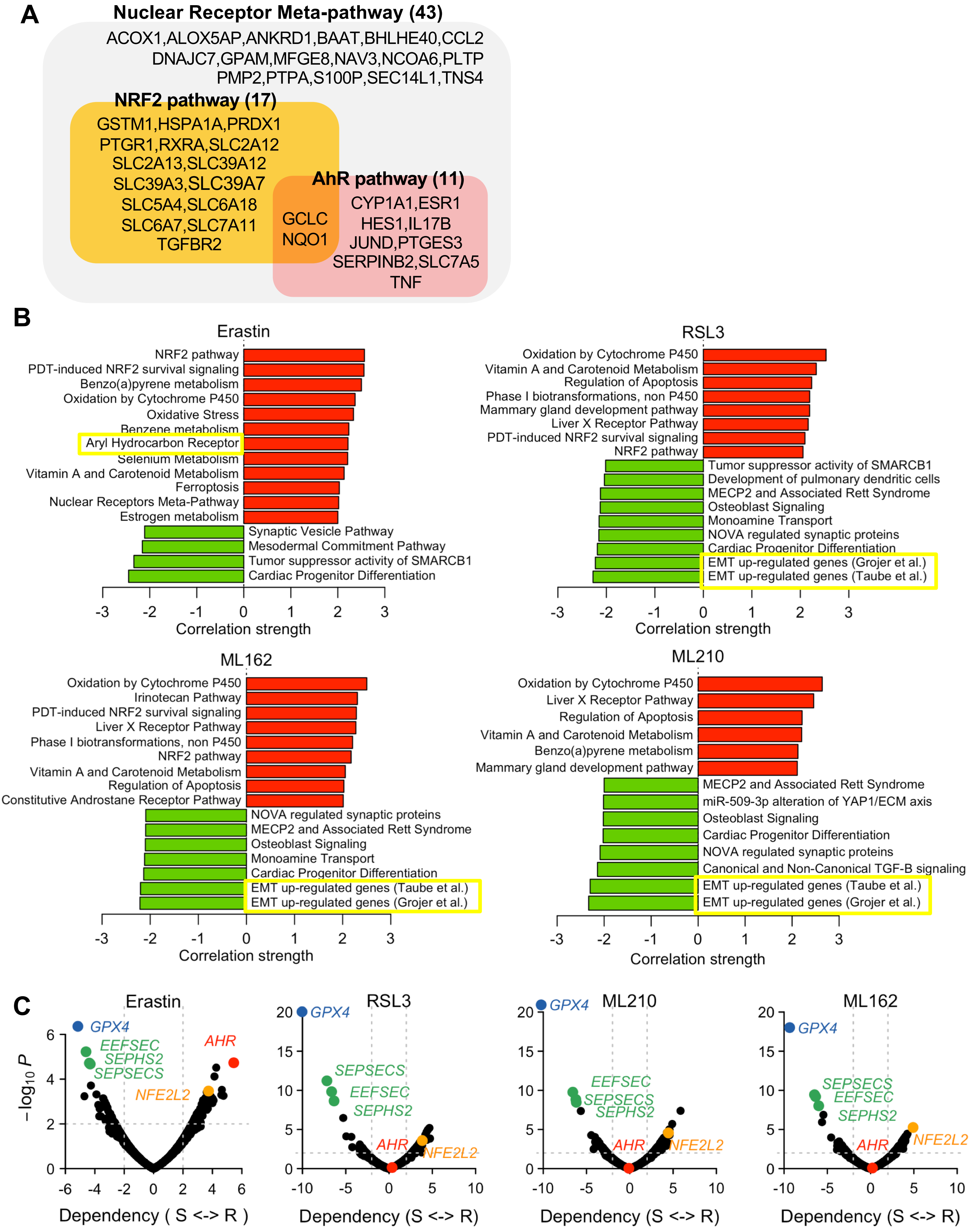
A
Nuclear Receptor Meta-pathway (43)
ACOX1,ALOX5AP,ANKRD1,BAAT,BHLHE40,CCL2
DNAJC7,GPAM,MFGE8,NAV3,NCOA6,PLTP
PMP2,PTPA,S100P,SEC14L1,TNS4
NRF2 pathway (17)
GSTM1,HSPA1A,PRDX1
PTGR1,RXRA,SLC2A12
SLC2A13,SLC39A12
SLC39A3,SLC39A7
SLC5A4,SLC6A18
SLC6A7,SLC7A11
TGFBR2
GCLC
NQO1
AhR pathway (11)
CYP1A1,ESR1
HES1,IL17B
JUND,PTGES3
SERPINB2,SLC7A5
TNF
B
Erastin
NRF2 pathway
PDT-induced NRF2 survival signaling
Benzo(a)pyrene metabolism
Oxidation by Cytochrome P450
Oxidative Stress
Benzene metabolism
Aryl Hydrocarbon Receptor
Selenium Metabolism
Vitamin A and Carotenoid Metabolism
Ferroptosis
Nuclear Receptors Meta-Pathway
Estrogen metabolism
Synaptic Vesicle Pathway
Mesodermal Commitment Pathway
Tumor suppressor activity of SMARCB1
Cardiac Progenitor Differentiation
-3 -2 -1 0 1 2 3
Correlation strength
RSL3
Oxidation by Cytochrome P450
Vitamin A and Carotenoid Metabolism
Regulation of Apoptosis
Phase I biotransformations, non P450
Mammary gland development pathway
Liver X Receptor Pathway
PDT-induced NRF2 survival signaling
NRF2 pathway
Tumor suppressor activity of SMARCB1
Development of pulmonary dendritic cells
MECP2 and Associated Rett Syndrome
Osteoblast Signaling
Monoamine Transport
NOVA regulated synaptic proteins
Cardiac Progenitor Differentiation
EMT up-regulated genes (Grojer et al.)
EMT up-regulated genes (Taube et al.)
-3 -2 -1 0 1 2 3
Correlation strength
ML162
Oxidation by Cytochrome P450
Irinotecan Pathway
PDT-induced NRF2 survival signaling
Liver X Receptor Pathway
Phase I biotransformations, non P450
NRF2 pathway
Vitamin A and Carotenoid Metabolism
Regulation of Apoptosis
Constitutive Androstane Receptor Pathway
NOVA regulated synaptic proteins
MECP2 and Associated Rett Syndrome
Osteoblast Signaling
Monoamine Transport
Cardiac Progenitor Differentiation
EMT up-regulated genes (Taube et al.)
EMT up-regulated genes (Grojer et al.)
-3 -2 -1 0 1 2 3
Correlation strength
ML210
Oxidation by Cytochrome P450
Liver X Receptor Pathway
Regulation of Apoptosis
Vitamin A and Carotenoid Metabolism
Benzo(a)pyrene metabolism
Mammary gland development pathway
MECP2 and Associated Rett Syndrome
miR-509-3p alteration of YAP1/ECM axis
Osteoblast Signaling
Cardiac Progenitor Differentiation
NOVA regulated synaptic proteins
Canonical and Non-Canonical TGF-B signaling
EMT up-regulated genes (Taube et al.)
EMT up-regulated genes (Grojer et al.)
-3 -2 -1 0 1 2 3
Correlation strength
C
Erastin
GPX4
EEFSEC
SEPHS2
SEPSECS
AHR
NFE2L2
−log10 P
0 2 4 6
-6 -4 -2 0 2 4 6
Dependency ( S <-> R )
RSL3
GPX4
SEPSECS
EEFSEC
SEPHS2
AHR
NFE2L2
0 5 10 15 20
-10 -5 0 5 10
Dependency (S <-> R)
ML210
GPX4
EEFSEC
SEPSECS
SEPHS2
AHR
NFE2L2
0 5 10 15 20
-10 -5 0 5 10
Dependency (S <-> R)
ML162
GPX4
SEPSECS
EEFSEC
SEPHS2
AHR
NFE2L2
0 5 10 15 20
-10 -5 0 5 10
Dependency (S <-> R)

**Figure S6**

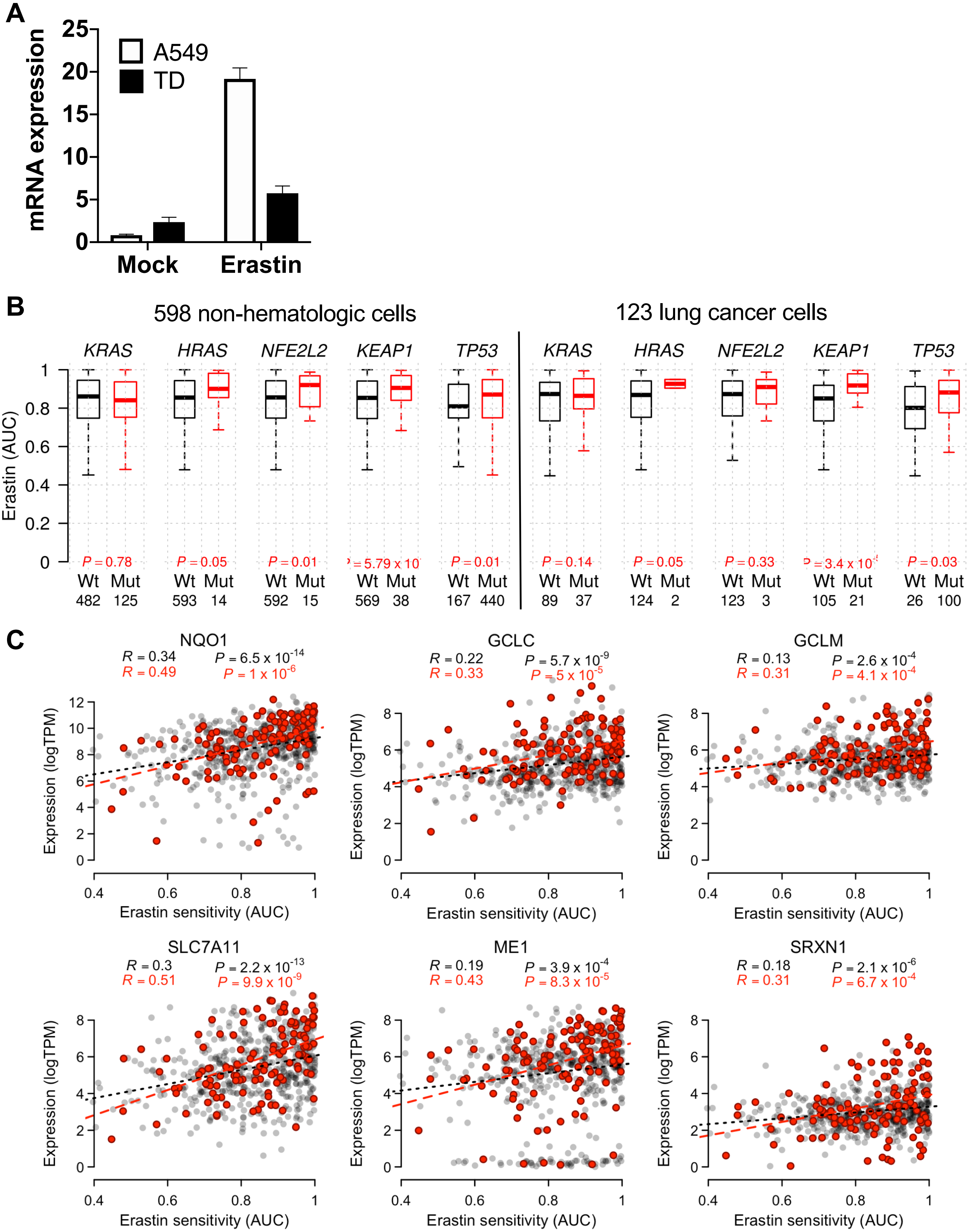
A
mRNA expression
A549
TD
Mock
Erastin
0
5
10
15
20
25
B
598 non-hematologic cells
123 lung cancer cells
KRAS
HRAS
NFE2L2
KEAP1
TP53
Erastin (AUC)
P = 0.78
P = 0.05
P = 0.01
= 5.79 x 10
P = 0.01
P = 0.14
P = 0.05
P = 0.33
= 3.4 x 10
P = 0.03
Wt Mut
482 125
593 14
592 15
569 38
167 440
89 37
124 2
123 3
105 21
26 100
C
NQO1
R = 0.34 P = 6.5 x 10-14
R = 0.49 P = 1 x 10-6
GCLC
R = 0.22 P = 5.7 x 10-9
R = 0.33 P = 5 x 10-5
GCLM
R = 0.13 P = 2.6 x 10-4
R = 0.31 P = 4.1 x 10-4
SLC7A11
R = 0.3 P = 2.2 x 10-13
R = 0.51 P = 9.9 x 10-9
ME1
R = 0.19 P = 3.9 x 10-4
R = 0.43 P = 8.3 x 10-5
SRXN1
R = 0.18 P = 2.1 x 10-6
R = 0.31 P = 6.7 x 10-4
Expression (logTPM)
Erastin sensitivity (AUC)

D

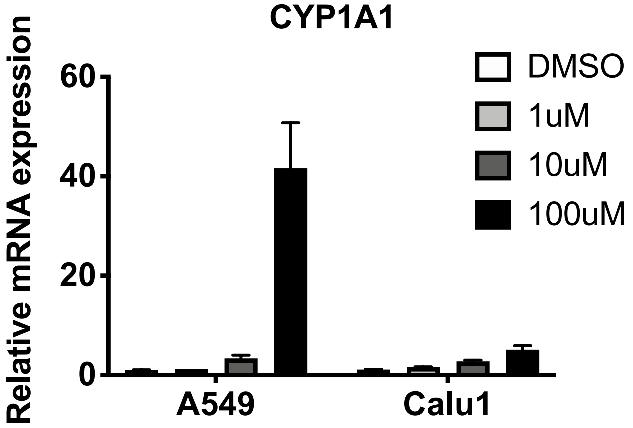
CYP1A1
Relative mRNA expression
60
40
20
0
A549
Calu1
DMSO
1uM
10uM
100uM

E

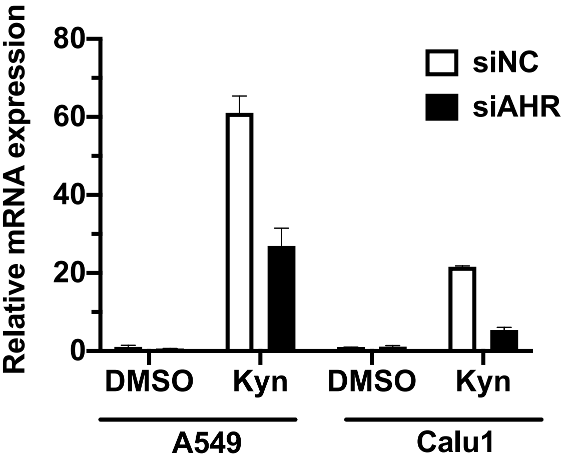
Relative mRNA expression
80
60
40
20
0
DMSO
Kyn
DMSO
Kyn
A549
Calu1
siNC
siAHR

### Figure S7

## A

| Cell line | Tissue | NRM prediction (AUC) | Group |
|---|---|---|---|
| A-704 | Kidney | 0.5144 | S |
| IMR-32 | Peripheral Nerve | 0.5674 | S |
| SK-N-MC | Primitive Neuroectoderm | 0.6240 | S |
| Calu-1 | Lung | 0.6666 | S.P.C |
| SW480 | Colorectal | 0.6832 | S |
| SNU-761 | Liver | 0.7092 | S |
| HCC33 | Lung | 0.7128 | S |
| HCC1359 | Lung | 0.7606 | R |
| NCI-H1299 | Lung | 0.7655 | R.P.C |
| A-549 | Lung | 0.9261 | R.P.C |
| NCI-H596 | Lung | 0.9944 | R |
| NCI-H1437 | Lung | 0.9947 | R |
